# Phenotypic plasticity and genetic control in colorectal cancer evolution

**DOI:** 10.1101/2021.07.18.451272

**Authors:** Jacob Househam, Timon Heide, George D Cresswell, Claire Lynn, Inmaculada Spiteri, Max Mossner, Chris Kimberley, Calum Gabbutt, Eszter Lakatos, Javier Fernandez-Mateos, Bingjie Chen, Luis Zapata, Chela James, Alison Berner, Melissa Schmidt, Ann-Marie Baker, Daniel Nichol, Helena Costa, Miriam Mitchinson, Marnix Jansen, Giulio Caravagna, Darryl Shibata, John Bridgewater, Manuel Rodriguez-Justo, Luca Magnani, Andrea Sottoriva, Trevor A Graham

## Abstract

Cancer evolution is driven by natural selection acting upon phenotypic trait variation. However, the extent to which phenotypic variation within a tumour is a consequence of intra-tumour genetic heterogeneity remains undetermined. Here we show that colorectal cancer cells frequently have highly plastic phenotypic traits *in vivo* in patient tumours. We measured the degree to which trait variation reflects genetic ancestry by quantifying the phylogenetic signal of gene expression across 297 samples with multi-region paired whole genome and transcriptome sequencing collected from 27 primary colorectal cancers. Within-tumour phylogenetic signal for genes and pathways was detected only infrequently, suggesting that the majority of intra-tumour variation in gene expression programmes was not strongly heritable. Expression quantitative trait loci analyses (eQTL) identified a small number of putative mechanisms of genetic control of gene expression due to the *cis*-acting coding, non-coding and structural genetic alteration, but most gene expression variation was not explained by our genetic analysis. Leveraging matched chromatin-accessibility sequencing data, enhancer mutations with *cis* regulatory effects on gene expression were associated with a change in chromatin accessibility, indicating that non-coding variation can have phenotypic consequence through modulation of the 3D architecture of the genome. This study maps the evolution of transcriptional variation during cancer evolution, highlighting that intra-tumour phenotypic plasticity is pervasive in colorectal malignancies, and may play key roles in further tumour evolution, from metastasis to therapy resistance.

## Introduction

Genetic intra-tumour heterogeneity (gITH) is an inevitable consequence of tumour evolution (Turajlic et al., 2019). Extensive gITH has been extensively documented across human cancer types (Black and McGranahan, 2021), and the precise pattern of gITH within an individual cancer is a direct consequence of the evolutionary dynamics driving the development of the tumour (Williams et al., 2019). Consequently, clones which experience positive, negative or neutral selection can be identified through analysis of gITH. However, natural selection in cancer operates on the phenotypic characteristics of a cell, for example the ability of a cancer cell to evade predation from the immune system (Rosenthal et al., 2019), or to metabolise in oxygen poor environments (Robertson-Tessi et al., 2015). Knowledge of the genotype-phenotype map of cancer cells is limited, and thus, whilst genomics offers us a window into determining *which* clones are selected, the methodology provides limited information on precisely *why* those clones are selected.

RNA sequencing (RNAseq) enables high throughput profiling of phenotypic characteristics of cancer cells by quantitative measurement of the gene expression levels that define the transcriptome (Stark et al., 2019). Historically, studies have focused on inter-tumour differences in gene expression patterns and have led to the identification of sets of genes with expression that is correlated with clinical outcomes; in colorectal cancer (CRC), the focus of this study, consensus molecular subtypes (CMS) (Guinney et al., 2015) or cancer cell intrinsic gene expression subtypes (CRIS) (Isella et al., 2017) exemplify this approach. Within these subtypes genotype-phenotype links have been noted, for instance the CMS1 subtype in CRC is highly enriched in tumours with mismatch repair deficiency, whereas *KRAS* mutations are enriched in CMS3 and the burden of somatic copy number alterations (SCNAs) is higher in CMS2&4 (Guinney et al., 2015; Lee et al., 2020), but the mechanistic relationship between the transcriptome and the genome remains to be determined. Furthermore, the growing popularity of multi-region sequencing, including single cell sequencing, has highlighted intra-tumour heterogeneity of the transcriptome (tITH) (for example in CRC see: (Lee et al., 2020; Roerink et al., 2018)). In CRC, multi-region sequencing studies have highlighted tITH of CMS and CRIS subtypes (Alderdice et al., 2018), and single cell sequencing shows cancer cells with differing levels of differentiation from a stem cell phenotype coexist within individual tumours (Lee et al., 2020).

Potentially tITH could be driven entirely by underlying (epi)genetic variation that evolves during tumour growth. However, the observation that local invasion is polyclonal in both CRC (Ryser et al., 2020) and in early breast cancer (Casasent et al., 2018), challenge the notion that cancer cell phenotype (here the ability to invade) is driven solely by the accrual of genetic mutations. Further, observations of rapid transcriptional shifts upon treatment (for example in melanoma (Shaffer et al., 2017)) and in CRC variation in subclone proliferation rates through serial re-transplantation rates despite largely stable patterns of genetic alterations (Kreso et al., 2013), discount the notion that transcriptomic phenotypes are determined solely by clonal replacement. Additionally, it has previously been determined that most driver mutations are clonal, meaning that this transcriptional variation must often happen in the absence of mutational drivers (Reiter et al., 2018). Collectively, these studies suggest that phenotypic characteristics are at least partially “plastic” – they can vary without requiring a new heritable (epi)genetic alteration to drive expression changes.

Here we perform multi-region paired transcriptomic and genomic profiling to characterise the evolution of phenotypic heterogeneity in colorectal cancers. We make use of the method of *phylogenetic signal* analysis developed in species evolution to determine the heritability of phenotypic traits from these data, and perform integrative analysis to elucidate candidate underlying molecular mechanisms controlling genetically-determined phenotypes.

## Results

We performed multi-region full-transcript RNA sequencing on 297 samples from 27 colorectal cancers (CRCs) (mean 11 samples/tumour, range: 1-38). The spatial sampling protocol and basic processing of these data are described in ref{epigenome}.

### Heterogeneity of gene and pathway expression in CRCs

We explored the heterogeneity of gene expression within and between CRCs. We selected genes that were frequently expressed in cancer cells by filtering for genes that were moderately-to-highly expressed (>=10 TPM) in at least 5% of tumour samples, and which did not show a significant negative correlation with sample purity (Methods). We clustered the filtered set of 11,401 genes using both the mean and variance of gene expression within each tumour (Figure 1A) and cut the dendrogram into four groups: group 1 had high average expression and relatively low variance in gene expression (“highly expressed, moderate heterogeneity”), groups 2 and 3 had progressively lower average gene expression and high variance in gene expression, while group 4 genes had low average gene expression and low variability between samples from the same tumour (Figure 1B&C). Meta-pathway analysis showed significant enrichment for pathways involved in cell growth and death in group 1, cancer-related genes in group 2 and pathways related to replication and repair in group 3. Group 4 was not significantly enriched for any class of pathways, and due to generally low expression and marked heterogeneity, was excluded from further analyses.

**Figure 1.**
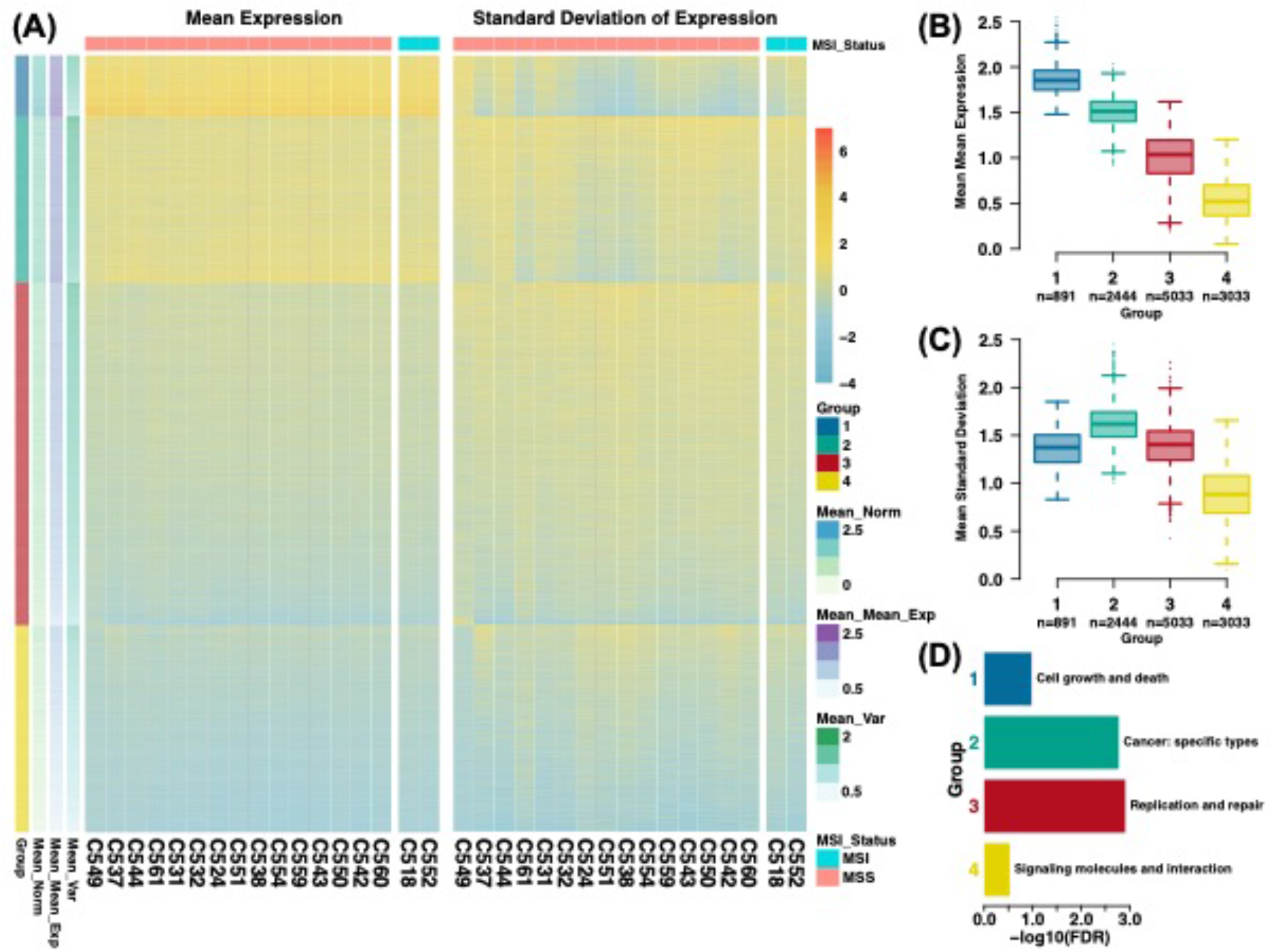
Clustering of genes based on mean tumour expression and intra-tumour heterogeneity of expression. **(A)** Heatmaps showing clustering of genes by expression level across tumours and expression variation within tumours. Hierarchical clustering revealed four distinct groups, named Group 1-4. Note units are scaled by column in each heatmap **(B)** Summary of mean expression level per Group. **(C)** Summary of intra-tumour heterogeneity of expression per Group, measured by standard deviation. **(D)** Meta-KEGG pathway analysis revealing which pathway categories are most over-represented in each Group (after removing “Infectious disease: bacterial” and “Neurodegenerative Disease” - most significant in Group 1).

We then repeated the clustering analysis using hallmark pathways (Liberzon et al., 2015) (Figure 2A) rather than individual genes cut the dendrogram into four groups of pathway enrichment. Group 1 contained pathways that were homogeneously enriched across all cancers, group 2 contained pathways with high average enrichment but high heterogeneity compared to group 1, and group 3 contained with lower average enrichment and heterogeneity compared to group 1. Group 4 contained only two pathways showed highly variable enrichment within each individual cancer (Figure 2B). Hallmark pathways were grouped into “classes” according to their biological mechanism (“oncogenic”, “immune”, “stromal”, etc) (Jiménez-Sánchez et al., 2020). Homogeneously enriched pathways showed moderate but not significant enrichment for cellular stress. Heterogeneously enriched pathways were not significantly enriched for any particular class of pathway (Figure 2D). Group 4 contained two pathways, epithelial-mesenchymal transition (EMT) and angiogenesis, and was enriched for “stromal” acting pathways.

**Figure 2.**
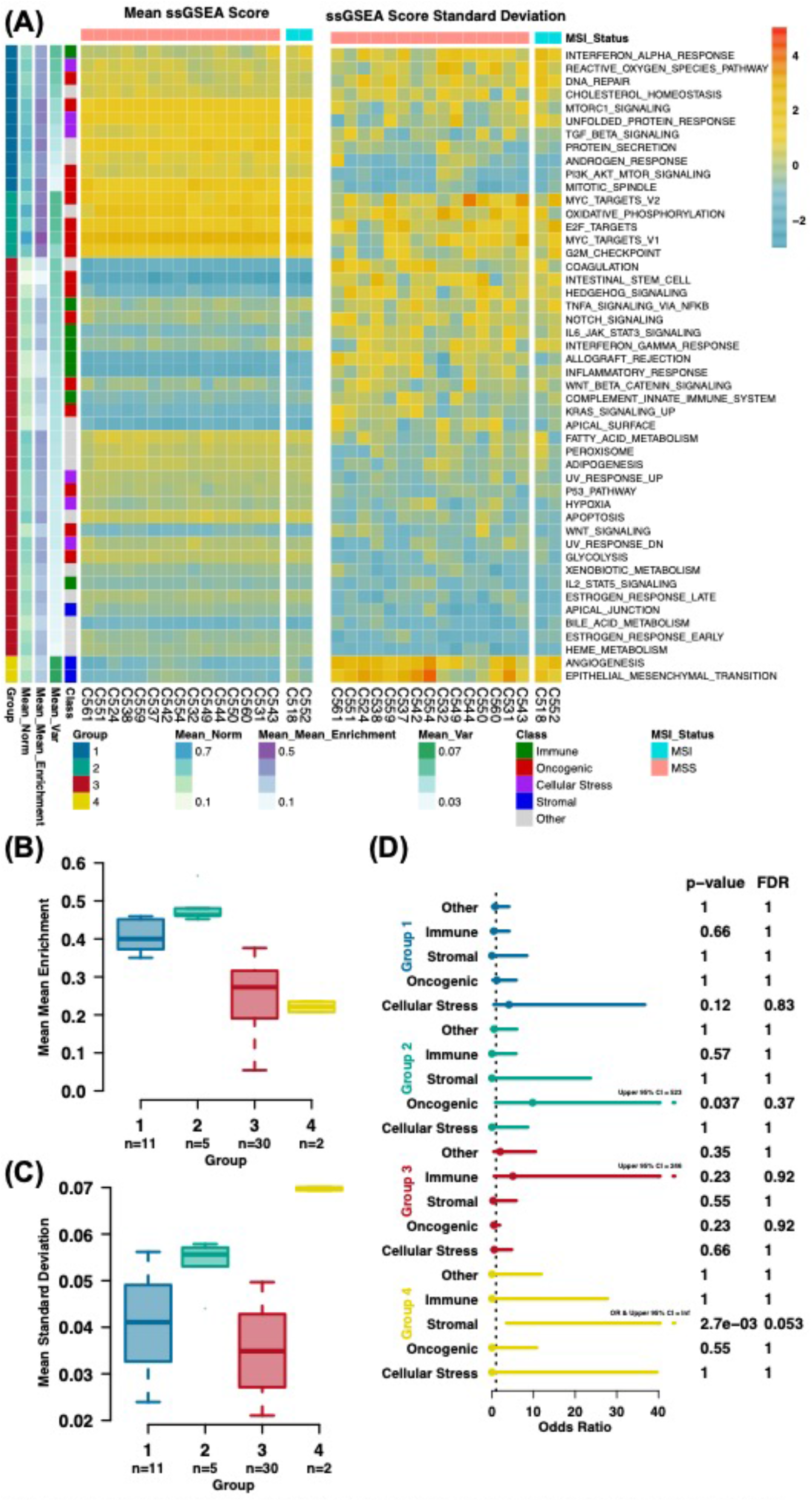
Filtering and clustering of pathways based on mean tumour enrichment and intra-tumour heterogeneity of enrichment. **(A)** Heatmaps showing clustering of pathways by enrichment level across tumours and enrichment variation within tumours. Hierarchical clustering revealed four distinct groups, named Group 1-4. Note units are scaled by column in both heatmaps **(B)** Summary of mean enrichment level per Class. **(C)** Summary of intra-tumour heterogeneity of enrichment per Class, measured by standard deviation. **(D)** Fisher’s exact test results comparing Groups to pathway classes.

Together, these analyses indicated that gene expression programs that define cancer cell biology and/or interaction with the surrounding immune microenvironment were not uniformly expressed across CRCs, despite the presumed importance of these phenotypes to cancer evolution.

Consensus molecular subtypes (CMS) and cancer cell intrinsic gene expression subtypes (CRIS) are systems to classify CRCs by gene expression patterns. We investigated the possibility of intra-tumour heterogeneity of these classifiers. 17 tumours had at least 5 tumour samples, and were amenable to these analyses. For CMS, only 2/17 tumours were homogeneously classified (both CMS3), 8 tumours contained samples assigned to two different classes, 3 tumours contained samples assigned to three classes while 4 tumours had samples assigned to all four CMS classes (Figure S1A). For CRIS only a single tumour was homogeneously classified (CRIS-A in C551), 3 tumours were assigned to 2 different CRIS classes, 10 were assigned to 3 classes and 3 assigned to 4 classes (Figure S1B). Overall, CRIS classification exhibited higher intra-tumour expression heterogeneity than CMS.

We assessed the consistency of assignment between CMS and CRIS: CMS3 classification correlated with CRISA classification, and CMS2 and CRISC classifications were also correlated, though many of these correlations were weak (Figure S1C). The genes used for both CMS and CRIS classification were found to be depleted in Group 1&2 genes (groups from Figure 1) and enriched in Group 4 genes (Figure S1D). Together these analyses indicate that both CRIS and CMS classifiers are sensitive to confounding by intra-tumour heterogeneity; inferring molecular subtype from a single biopsy risk incorrect classification of the tumour as a whole. Potentially gene expression classifiers robust to intra-tumour heterogeneity could be constructed by limiting included genes to those found only in Group 1 genes (those with high expression and low intra-tumour heterogeneity) analogous to the approach in (Biswas et al., 2019).

### Evolutionary dynamics of gene and pathway expression heterogeneity

We sought to understand the evolutionary dynamics of the observed heterogeneous patterns of gene expression. Paired whole genome sequencing data (either deep sequencing [dWGS] or shallow sequencing [sWGS] data that enabled genotyping) was available for 157/279 samples with matching RNAseq data. We constructed phylogenetic trees that depicted the shared genetic ancestry of the set of samples from each tumour (method described in associated paper EPIGENOME) and excluded tumours with fewer than 6 paired DNA-RNA samples (leaving 114 samples from 8 tumours, median 11 samples per tumour, range 6-31). The terminal nodes of the trees were the extant samples – i.e. the tumour subclones that we had phenotypically characterised with RNAseq data - and so we overlaid the measured gene expression profiles onto the phylogenetic trees (Figure 3C&D and Figure S2).

**Figure 3.**
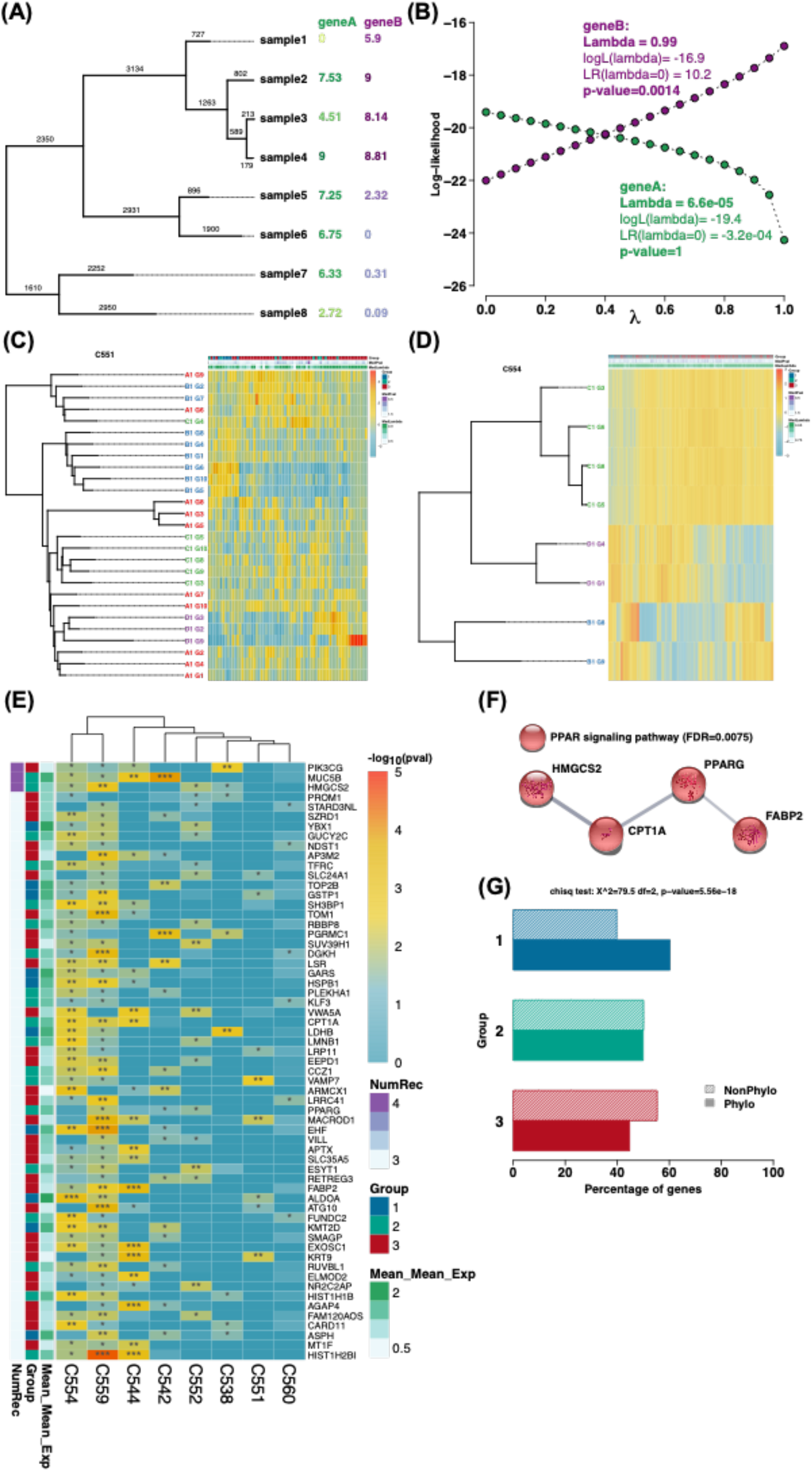
Measuring of phylogenetic signal of gene expression with Pagel’s lambda **(A)** Example tree with branch lengths and expression of two example genes shown. GeneA’s expression was randomly generated for each sample, while GeneB’s expression is from a simulation of Brownian Motion (BM) along the tree, meaning expression will be more similar for closely related samples. Expression for both genes is scaled between 0 and 9. **(B)** Log-likelihood values are calculated for all values of lambda between 0 and 1, where a lambda of 1 means the structure of the tree itself explains the gene expression while a lambda of 0 means that the tree has to lose all of its structure to explain the evolution of the gene expression under BM. The lambda estimate is the lambda with the maximum log-likelihood and a likelihood ratio test against lambda=0 then tests for the significance of phylogenetic signal. **(C)** and **(D)** Phylogenetic trees and heatmaps of genes with significantly high phylogenetic signal for tumours C552 and C554 respectively. **(E)** Genes with recurrent phylogenetic signal across tumours, genes shown were found to have significantly high phylogenetic signal in at least three tumours **(F)** Enrichment of KEGG ‘PPAR signaling pathway’ for recurrently phylogenetic genes. **(G)** Results of chi-squared test showing whether gene groups were enriched for phylogenetic genes.

Phylogenetic signal is a statistical method from evolutionary biology that measures the degree to which the phenotypic (dis)similarity between species is explained by genetic ancestry, and is quantified by the Pagel’s lambda statistic (Freckleton et al., 2002; Pagel, 1999) (Figure 3A&B). We assessed the phylogenetic signal of gene expression heterogeneity in each of our CRCs with sufficient paired RNA-seq-WGS data. Pagel’s lambda was computed for 8368 genes from groups 1- 3 (as defined in Figure 1). We identified significant phylogenetic signal by comparing to the null hypothesis of no phylogenetic signal (gene expression unrelated to genetic ancestry) using a likelihood ratio test (see explanatory Figure 3A&B). Figure 3C&D show the expression of significantly phylogenetic genes mapped onto phylogenetic trees for two individual tumours (see Figure S2 for all tumours used).

A median of 166 genes had phylogenetic signal (p<0.05) within each tumour (range 67-2335), and the number of genes with phylogenetic signal did not significantly correlate with the number of samples per tumour (p=0.25, Figure S3). Group 1 genes (highly expressed, moderate heterogeneity) were enriched for phylogenetic signal, whereas group 3 genes (moderately expressed, moderate heterogeneity) were significantly depleted for phylogenetic signal (Figure 3G).

We searched for genes that recurrently showed phylogenetic signal across tumours. 52 genes were found to be phylogenetic in at least three tumours (Figure 3E). The KEGG pathway ‘PPAR signalling’ (peroxisome proliferator-activated receptor signalling) that is involved in prostaglandin and fatty acid metabolism (Michalik et al., 2004) was statistically over-represented in this gene set (FDR=0.0075; Figure 3F; stringDB analysis). Links between PPAR metabolism and CRC have been previously reported (Currie et al., 2013; Fuchs et al., 2005).

We then assessed phylogenetic signal at the level of gene expression pathways. Figure 4A&B depicts phylogenetic signal for gene expression pathways for tumours C554 and C559 (see Figure S5 for all eight tumours analysed). Two pathways were recurrently phylogenetic in at least 3 tumours: fatty acid metabolism, related to PPAR signalling which was identified in the gene-level analysis, and ‘MYC_TARGETS_V2’ that contains genes regulated by *MYC* signalling (Figure 4B). 23 pathways were significantly phylogenetic in at least 2 tumours, and no recurrently phylogenetic pathways were consistently associated with the pathway expression-heterogeneity subgroups identified in Figure 2 (Figure 4C). Thus, with the exception of MYC signalling and PPAR fatty acid metabolism, the gene and hallmark pathway expression levels only infrequently showed strongly heritable subclonal variation during CRC evolution.

**Figure 4.**
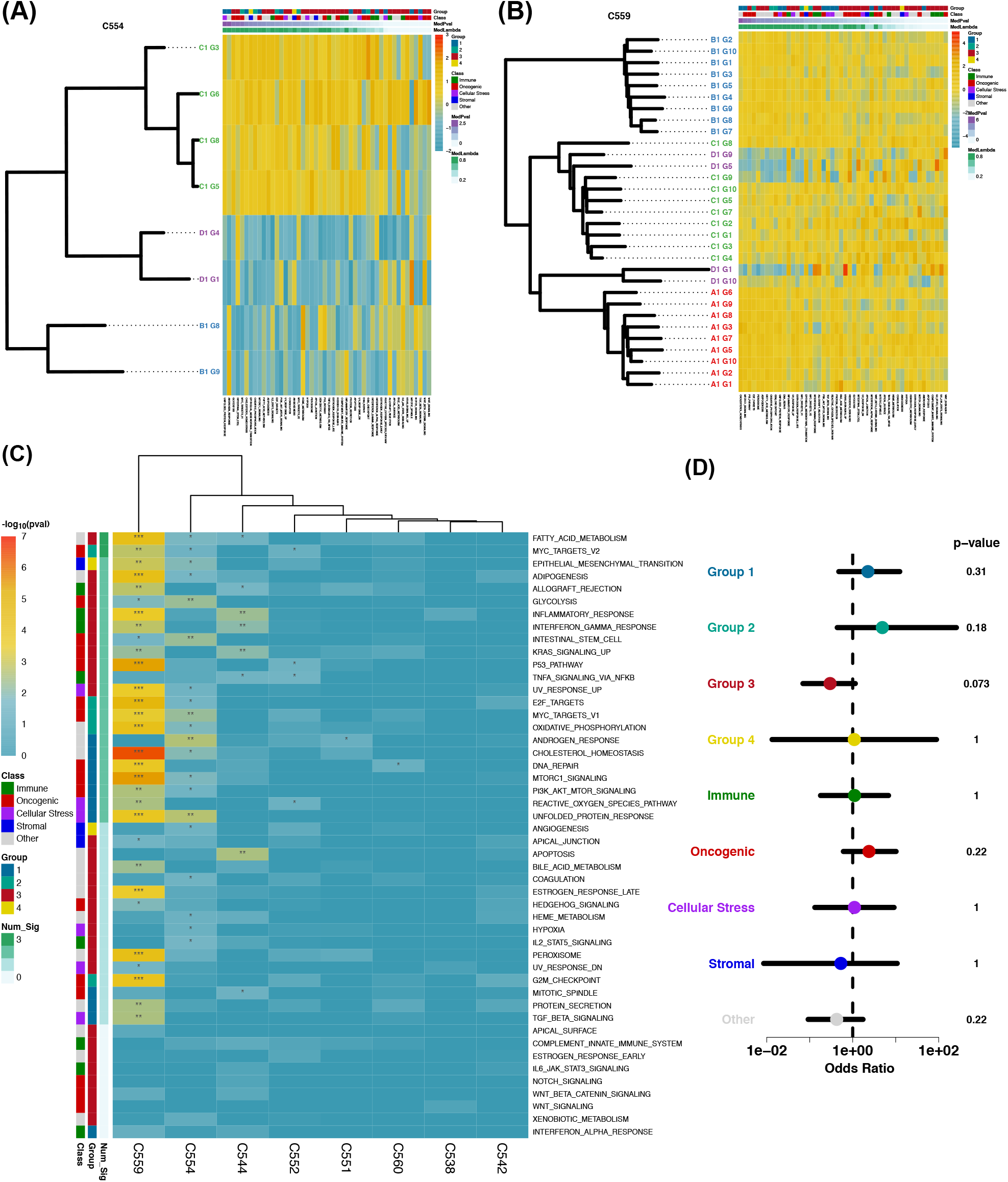
Phylogenetic signal of pathways. **(A)** and **(B)** Example phylogenetic trees and pathway enrichment heatmaps for tumours C554 and C559 respectively. Pathways are ordered by decreasing phylogenetic signal. **(C)** Heatmap showing recurrence of phylogenetic signal of pathways across tumours. Pathways are ordered by decreasing recurrence. **(D)** Results of fisher tests investigating whether pathways that are recurrently phylogenetic (in at least 2 tumours) are enriched for a particular group or class.

We assessed the power of our analysis to detect phylogenetic signals given the size and structure of our dataset (a *post hoc* power calculation). To do this we performed simulations of gene expression evolution across the phylogenetic trees observed in our cohort CRCs, where gene expression was Poisson distributed across nodes and was increased by a factor of 5-100% in a randomly chosen clade of the tree. We had 90% power to detect heritable changes in gene expression bigger than 85%, which occurred sufficiently early, in 87.5% (7/8) of the tumours in our cohort (Figure S4). We note that only one tumour (C559) had phylogenetic signals that were significant after multiple testing correction; our power analysis indicated that our power to detect associations was greatest for this tumour (90% power to detect appropriately-timed heritable changes in gene expression >20%). Thus, power analysis indicated that our dataset was sufficient to enable detection of subclonal, large-effect and heritable changes in gene expression, and these events were rare within our cohort of CRCs.

Together, the gene and pathway level assessments of the evolution of transcriptional heterogeneity show only few occurrences of substantial subclonal changes in transcription activity. In other words, transcriptional activity tends to be somewhat ‘uniform’ across CRCs. Natural selection acts upon phenotypic variation, and so commonplace phenotypic uniformity is consistent with prior reports that analysed genetic data and found only infrequent evidence of stringent subclonal selection in CRCs (Sottoriva et al., 2015; Sun et al., 2017; Williams et al., 2018).

### Genetic determinants of gene expression heterogeneity

Genes that recurrently showed phylogenetic signal across multiple patients in our dataset were rare, but 4006/8368 (47.9%) of genes showed some evidence of heritable changes in gene expression in a single cancer. Somatic mutations altering gene expression are a potential mechanistic explanation of phylogenetic signal. We used a regression framework (see Methods), analogous to expression quantitative trait loci (eQTL) used in human population genetics(Nica and Dermitzakis, 2013), to detect significant *cis*-associations between inter- and intra-tumour somatic genetic heterogeneity (see Methods) and gene expression.

5,927 genes had *cis-* somatic genetic variation (a genic or non-genic (promoter or enhancer) somatic SNV) in at least two samples across our cohort (out of 167 samples collected from 19 tumours where there were at least two samples with matched RNAseq and dWGS). The association between the expression of each of these genes and the somatic genetic alterations was examined using multivariate linear regression on z-score normalised expression values, revealing 1,402 genes (FDR<0.01) with significant associations (Figure 5), which we termed “eQTL gnenes”.

**Figure 5.**
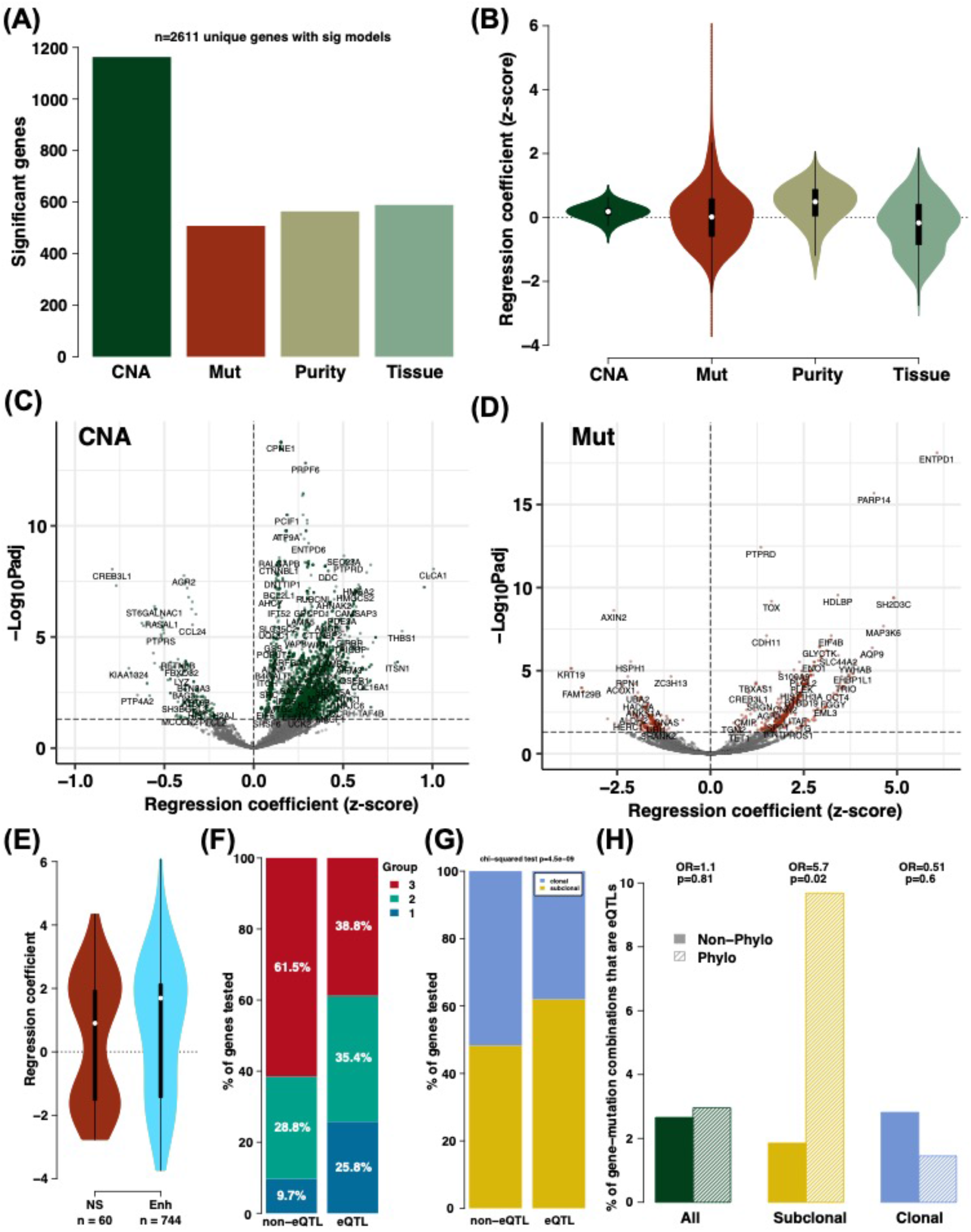
eQTL analysis. **(A)** The number of genes with significant models for each data type. **(B)** The distribution of regression coefficients (effect sizes) for each data type. **(C)** and **(D)** Volcano plots highlighting selected genes that were significant for CNA and Mut eQTLs respectively. **(E)** In comparison to non-synonymous SNVs (NS), enhancer (Enh) mutations tended to have large effect sizes and a higher proportion of positive effect sizes. **(F)** eQTL associated genes were enriched for Group 1 and 2 genes and depleted in Group 3 genes in comparison to non-eQTL associated genes. **(G)** The proportion of eQTL associated genes that were subclonal was higher than the same proportion for non-eQTL associated genes. **(H)** Visualisation of Fisher’s exact tests showing that gene-mutation combinations were more likely to be eQTLs if they were associated with recurrent phylogenetic genes (genes found to be phylogenetic in at least 3 tumours) for subclonal mutations and that this remains true, but not significant, when considering all mutations, but not for clonal mutations.

Of these 1,402 eQTL genes, SCNAs contributed to expression changes of 1163/1402 (83.0%) genes (Figure 5C; Table S1), but the magnitude of the effect on expression tended to be small (median effect size 0.30 standard deviation in expression change per allele copy). A positive correlation between copy number and expression was observed for 1082 genes, but interestingly a negative correlation was observed for 81 genes. Amongst the associations, genetic deletions were enriched at genes where genetic copy number was positively associated with expression (i.e. genes where genetic deletions cause a decrease in gene expression or genetic gains increase expression; p=5.9e-05; Figure S6A), whereas genes with negative associations with copy number (i.e. genic deletions associated with an increase in gene expression, or genetic gains decreases in expression) were enriched for loci with copy number 3 (p=6.7e-06; Figure S6C, unbalanced gain) but not 4 (p=0.033; Figure S6D, commonly balanced gain). Consequently, we speculate that this is due to dominant-negative activity of the amplified allele.

Somatic SNVs, either coding or non-coding variants, were associated with gene expression variation in 508 genes (Figure 5D; Table S1), and typically the magnitude of the association was much greater than for CNAs (mean effect size 1.92 vs 0.30 standard deviations for SNV vs single copy number change, Figure 5B). For coding somatic mutations, approximately equal numbers of associations that were associated with an increase versus decrease in expression were observed (33 coding SNVs increase expression versus 27 decreasing expression; p=0.4). Non-coding enhancer somatic mutations were associated with the largest changes in gene expression observed in our cohort, and were more likely to increase expression (486 increase vs 258 decrease; Figure 5E, p=6.3e−^17^). The expression of 175 genes was significantly associated with both CNAs and SNVs, indicating how the combination of somatic mutation and copy number alterations together determine the gene expression phenotype of cancer cells. We also found that the proportions of genes assigned to the expression groups identified in Figure 1 were significantly different between non-eQTL associated genes and eQTL associated genes (Figure 5F; chi-squared test, p-value=1.76e^−36^). Specifically, eQTL associated genes were enriched for Group 1&2 genes, while they were depleted for Group 3 genes. This is most likely due to the fact that we had the most power to detect mutation-expression associations in genes with relatively high expression, and Group 1&2 genes had higher mean expression than Group 3 genes (see Figure 1B).

Leveraging our multi-region sequencing data, we assessed the clonality of mutations that were associated with differential gene expression. Subclonal mutations were slightly but significantly more likely to be associated with *cis* gene expression changes than clonal mutations (Figure 5E; X^2^=34.4, p=4.5e-09, Figure 5G), indicating that “*trans*” effects due to the (epi)genetic state of a cancer cell commonly determine whether or not a somatic mutation will cause changes in gene expression. Furthermore, we calculated how often somatic mutations explained the observed phylogenetic signal (i.e. from Figure 3). Subclonal mutations that associated with changes in *cis* gene expression were enriched for phylogenetic signal (OR=5.7, p=0.02; Figure 5G), and this enrichment was absent when examining all mutations (OR=1.1, p=0.81) or clonal mutations (OR=0.51, p=0.6), see Figure 5H. Thus, collectively our data show that subclonal mutations within cancers are associated with frequent and heritable changes in *cis* gene expression.

We used data from N=394 metastatic colorectal cancers from the Hartwig cohort {31645765} where there was paired dWGS and RNA-seq data available to validate the *cis* effects of somatic mutations on gene expression. 115/508 “*cis*-acting mutations” identified in our dataset were present in the Hartwig dataset in more than 1 sample, and the correlation between mutation and gene expression observed in our cohort was detected in Hartwig samples for 9 of these 115 mutations (Figure S7A-I). For 93/ 106 mutations that did not validate, *post hoc* power assessment indicated insufficient power (too few cancers with the mutation) in the Hartwig cohort. For the remaining 13 mutations, we had observed the *cis* effect on gene expression only when the mutation acted sub-clonally within a tumour for 2/13 mutations. For the remaining 11/13 mutations, we assume that failure to validate expression effects in Hartwig is due to unexplained “*trans*” effects (Figure S7J).

We tested if subclonal eQTL mutations were associated with subclonal expansions by assessing the sizes of clades on phylogenetic trees that carried the eQTL variants. Population genetic theory predicts that clade sizes for neutral variants are inversely proportional to clade age, which can be proxy-measured by the number of mutations accumulated since the most recent common ancestor (MRCA) of the clade (method is described in accompanying manuscript INFERENCE). Examination of the relationship between clone size against the time to MRCA revealed that the clades with subclonal eQTL mutations were broadly consistent with clade sizes expected from neutral mutations (Figure S8). This demonstrated that, in cases where gene expression is genetically determined, these expression changes were not likely to be experiencing strong clonal selection, and so were likely effectively-neutral.

### Non-coding somatic mutations are associated with changed chromatin accessibility and differential gene expression

We sought to identify the mechanism of how non-coding enhancer somatic mutations cause changes in *cis* gene expression. We leveraged paired chromatin accessibility sequencing (ATACseq) data (n=247 with matched mutation and ATACseq data, n=108 with matched mutation, RNA and ATACseq data) to explore changes in chromatin architecture associated with enhancer somatic mutations (Figure 6 & Figure S9).

**Figure 6.**
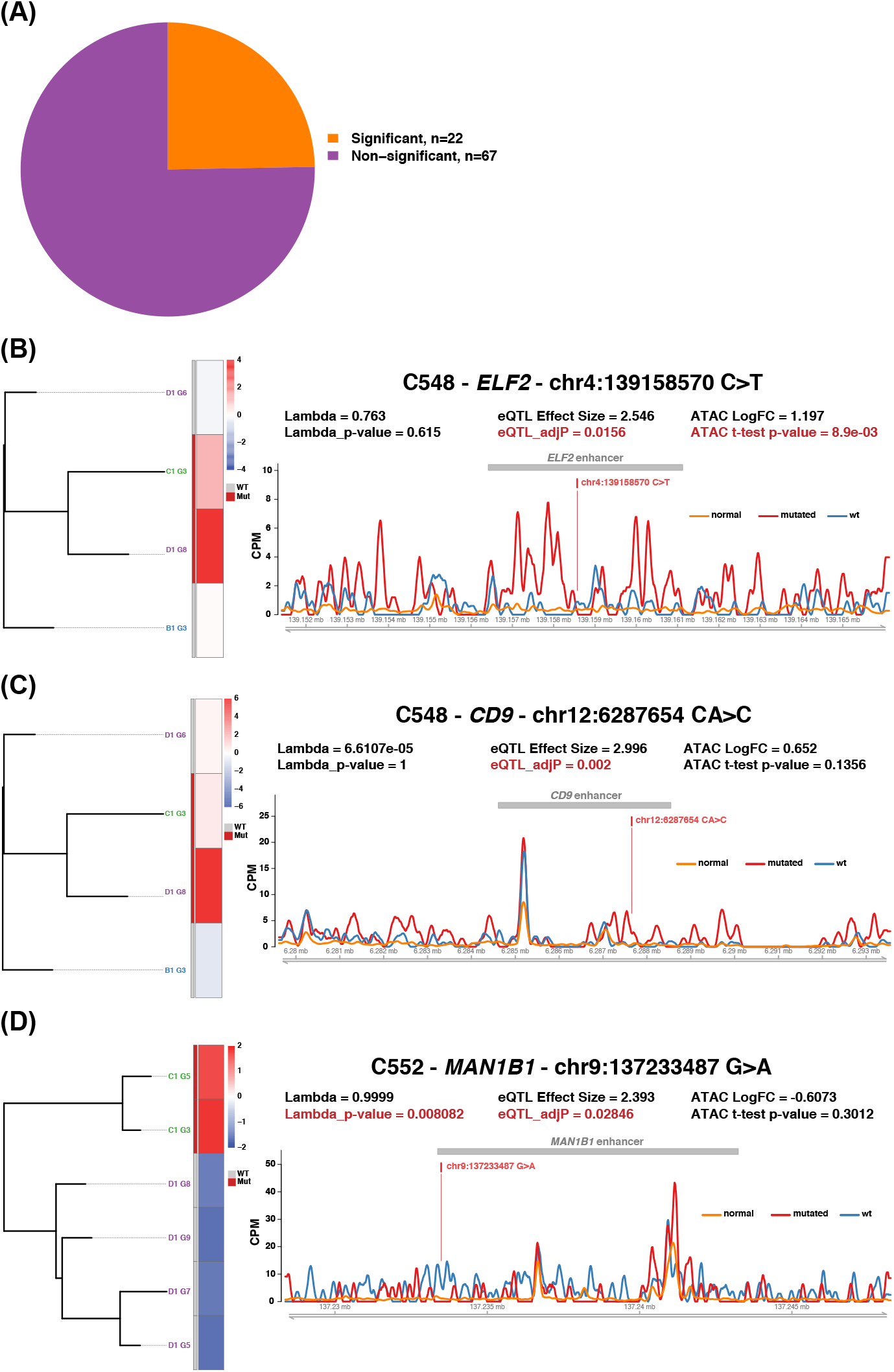
Combining eQTLs ATAC-seq and phylogenetic data. **(A)** Pie chart showing the proportion of subclonal enhancer eQTLs that displayed significant changes in chromatin accessibility via ATAC-seq **(B)** An example of an eQTL with supporting ATAC-seq data - ELF2 in tumour C548. **(C)** An example of an eQTL with limited supporting ATAC-seq data - CD9 in tumour C548. **(D)** An example of a phylogenetic eQTL with limited supporting ATAC-seq data - MAN1B1 in tumour C552.

22/89 enhancer mutations were associated with altered chromatin accessibility within cancerous tissue (Figure 6A). Somatic enhancer mutations associated with increased gene expression were typically associated with more open chromatin and vice versa (Figure 6B-D and Figure S9). We note that many enhancer mutations had no discernible effect on the ATACseq signal, highlighting that biological consequences of enhancer mutations can occur through a variety of mechanisms not necessarily affecting over accessibility of the DNA.

Striking examples of putatively mutation-induced changes in chromatin architecture included somatic mutations in the enhancers of the transcription factor *ELF2* (an effector of MYC signalling (Schmidt et al., 2019)), *CD9* (a tetraspanin member whose increased expression weakly correlates survival in CRCs (Kim et al., 2016) and *MAN1B1* (a mannosidase implicated in protein folding (Sun et al., 2020)). All three were associated with increased expression and visibly increased chromatin accessibility (Figure 6B-D), and *MAN1B1* was also determined to have significant phylogenetic signal. The mutation in *MAN1B1* is also noticeably displaced from the main affected peak of ATACseq signal, suggesting that broadly altered chromatin accessibility of the locus rather than a binary switch between “open” and “closed” states. We note that some enhancer mutations were found to decrease both expression and chromatin accessibility (see Figure S9A&B).

## Discussion

Heterogeneity in gene expression is common between and within colorectal cancers. Leveraging the fact that clone ancestry is encoded by somatic mutations in the genome, here we determined that only a small proportion of the observed transcriptomic variation shows evidence of recurrent heritability through tumour evolution (<1% of expressed genes and <5% of hallmark pathways), with a maximum of 28% of genes showing heritable expression in any individual tumour. This points towards phenotypic plasticity – the ability of a cancer cell to change phenotype without underlying heritable (epi)genetic change – to be a common phenomena in CRC. We previously hypothesised that the observation of infrequent stringent selection for subclones within CRCs was consistent with the notion that phenotypic plasticity was established within cancer cells at the outset of cancer growth (Sottoriva et al., 2015). Here our explicit analysis of transcriptomic variation confirms this hypothesis.

Nevertheless, we find that somatic mutations do, infrequently, cause detectable and heritable changes in gene expression. Of 29,949 associations between somatic mutations and gene expression examined here, only 796 were associated with significant changes in *cis* gene expression and so can be thought of as “functional” mutations. Moreover, subclonal functional mutations were rare: in any individual tumour, we detected a median of 1 (max 34) putatively functional subclonal mutations and so the vast majority of genes had expression that was not controlled by new somatic mutations which had arisen during tumour evolution.

We note that phenotypic changes do not necessarily correlate with changes in fitness – the newly induced expression of a particular gene may have no relevance to the ability of that cell to survive or grow in its current microenvironment, and indeed across species most genetic “tinkering” is near-neutral or even deleterious (Eyre-Walker and Keightley, 2007). In cancers, genetic analysis predicts the vast majority of SNVs to be neutral (Martincorena et al., 2017). Indeed, our related analysis shows that tumour subclones which have, or have previously had, increased fitness show at most only slight transcriptomic differences within the extant tumour (reference associated INFERENCE manuscript). Thus, at least some of the observed tITH is simply “noise” produced by the stochastic accumulation of mutations during tumour growth, and care should be taken not to conflate transcriptional variation with evidence of important variation in tumour cell biology.

Our study shows the evolutionary origins of gene expression heterogeneity in cancer can be mechanistically studied by combining genomics with transcriptomics through the lens of evolutionary biology.

## Methods

### Sample preparation and sequencing

The method of sample collection and processing is described in an accompany manuscript (ref associated PROTOCOL manuscript). Sequencing and basic bioinformatic processing of DNA-sequencing and ATAC-seq data are described in a second accompanying manuscript (ref associated EPIGENOME manuscript).

### Processing of RNA-seq

After initial quality control with FastQC (Andrews, 2021) and default adaptor trimming with Skewer (Jiang et al., 2014), paired-end reads were aligned to GRCh38 reference genome and version 28 of the Gencode GTF annotation using the STAR 2-pass method (Dobin et al., 2013). Read groups were added with Picard v.2.5.0 {http://broadinstitute.github.io/picard}. Per gene read counts were produced with htseq-count that is incorporated into the STAR pipeline (Anders et al., 2015).

### Sample filtering

Raw gene counts were first filtered for reads uniquely assigned to non-ribosomal protein-coding genes located on canonical chromosomes (chr1-22, X and Y).

If samples had less than 5M of these ‘usable’ reads they were re-sequenced to improve coverage. Where possible, the same library preparation pool was sent again for sequencing. These `top-ups' proved to be true technical replicates, since the resulting gene expression of the re-sequenced samples clustered very closely to their original samples on both a sample-sample heatmap and a principal component analysis (PCA). It was therefore determined that the fastqs of these samples could simply be merged at the start of the pipeline. In cases where resequencing was required but insufficient library remained, a new library was prepared and the sequencing run that produced the highest read was used in subsequent analysis. For 8 samples, the sequencing of the second library contained too few reads to enable downstream analysis. 6/8 samples showed per gene read counts that were very similar between libraries 1 and 2 (Spearman’s rank correlation between replicates was significantly higher than the mean; Wilcoxon one-way rank test; FDR<0.01) and so reads were combined across libraries, the other 2/8 samples were discarded.

Samples were also discarded if matched DNA-sequencing revealed a tumour purity of less than 0.05.

### Gene expression normalisation and filtering

The number of non-ribosomal protein coding genes on the 23 canonical chromosome pairs used for quality control was 19,671. Raw read counts uniquely assigned to these genes were converted into both transcripts per million (TPM) and variance-stabilising transformed (vst) counts via DESeq2 (Love et al., 2014).

A list of expressed genes (n=11,667) was determined by filtering out genes for which less than 5% of tumour samples had at least 10TPM. In order to concentrate on tumour epithelial cell gene expression, genes were further filtered out if they negatively correlated with purity as estimated from matched DNA sequencing data (see associated manuscript EPIGENOME for methodology of purity estimation). Specifically, for the 157 tumour samples that had matched DNA-sequencing and therefore accurate purity estimates, a linear mixed effects model of Exp (vst) ~ Purity + (1|Patient) was compared via a chi-squared test to Exp ~ (1|Patient). Genes which had a negative coefficient for Purity in the first model and an FDR adjusted p-value less than 0.05, suggesting that Purity significantly affected the expression, were filtered out. This led to a filtered list of 11,401 expressed genes.

### Gene expression clustering

For each tumour with at least 5 tumour samples (n=17 tumours), mean expression and standard deviation of expression was calculated for every filtered expressed gene (n=11,401) using DESeq2’s vst normalised counts. Euclidean distance matrices of mean expression and standard deviation of expression were calculated based on non-MSI tumours. Distance matrices were combined with ‘fuse’ from the ‘analogue’ R package https://cran.r-project.org/package=analogue with equal (50:50) weighting and complete linkage hierarchical clustering was performed. 4 gene groups were determined using cutree(k=4). For plotting of Figure 1A tumours were clustered with the same approach as above and both mean expression and standard deviation of expression matrices were scaled by columns.

Conversion to entrez gene IDs and gene symbols was carried out using biomaRt {ref}, using Ensembl version 90. Where IDs were missing, newer Ensembl versions and manual curation was used, the full list of gene information is available (Table S2).

For the KEGG meta-pathway analysis, pathways and pathway categories were downloaded from https://www.kegg.jp/kegg-bin/show_brite?hsa00001_drug.keg. The enrichment of KEGG pathways for each gene group was determined with enrichKEGG from clusterProfiler (Yu et al., 2012), and pathways enriched at FDR<0.1 were inputted into ‘enricher’ to determine pathway category enrichment (FDR<0.1). Pathway categories "Neurodegenerative disease" and "Infectious disease: bacterial" were removed due to irrelevance to colorectal cancer cell biology.

### Pathway enrichment clustering

Hallmark pathways were download from MSigDB (Liberzon et al., 2015) and un-related pathways (SPERMATOGENSIS, MYOGENESIS and PANCREAS_BETA_CELLS) were removed from analysis while the COMPLEMENT pathway was renamed to COMPLEMENT_INNATE_IMMUNE_SYSTEM. Additional INTESTINAL_STEM_CELL {21419747} and WNT_SIGNALING {http://www.gsea-msigdb.org/gsea/msigdb/geneset_page.jsp?geneSetName=WNT_SIGNALING} pathways were added.

For each multi-region tumour (n=17), the TPM expression of protein-coding genes converted to entrez gene IDs (n=18,950) was used as input for single sample gene set enrichment analysis using the GSVA R package (Hänzelmann et al., 2013). The mean and standard deviation of enrichment was then recorded for each tumour. KRAS_SIGNALING_DN had average enrichment below zero so was removed from downstream analysis, leading to a final list of 48 pathways.

Analogous to the genic analysis, mean and standard deviation of pathway enrichment were jointly used to determine 4 groups of pathways while tumours were clustered and matrices normalised by column as before. Fisher’s exact tests were subsequently performed to determine if pathway classes (Jiménez-Sánchez et al., 2020) were significantly enriched/depleted in particular pathway groups.

CMS and CRIS classifications were determined using the CMScaller R package (Eide et al., 2017). As recommended, raw gene counts were used as input with ‘RNAseq=TRUE’, meaning these counts underwent log2-transformation and quantile normalisation. CMS and CRIS were predicted using the templates provided in the CMScaller package and samples were assigned to the subtype with the shortest distance.

### Phylogenetic signal analysis

Phylogenetic trees were determined as described in methods of {inference paper} with shallow WGS (sWGS) samples genotyped onto deep WGS (dWGS) trees. Tumours with fewer than 6 paired DNA-RNA samples were excluded from this analysis leaving 114 samples from 8 tumours (median 11 samples per tumour, range 6-31).

Added sWGS samples, however, had zero branch lengths as mutations unique to a sample could not be called with sWGS methodology. To account for these “missing” unique variants, we inferred the likely number of unique variants from the matched dWGS samples. For each sWGS sample from a particular tumour region, a new tip branch length (“leaf length”) was drawn from a Poisson distribution based on the mean number of unique mutations observed in each dWGS sample from the same spatial tumour region. DNA samples that did not have matched RNA-seq samples were then removed from the trees (with drop.tip from ape R package (Paradis and Schliep, 2019). This was process was repeated 100 times for each tumour, leading to a forest of 100 phylogenetic trees with slightly varying branch lengths for each sWGS sample.

In the genic phylogenetic signal analysis, Pagel’s lambda was calculated for Group 1-3 genes (n=8,368) using phylosig from the phytools R package (Revell, 2012). This returns the maximum likelihood Pagel’s lambda estimate and a p-value for the likelihood ratio test with the null hypothesis of lambda=0. This analysis was performed for all 100 trees and the median lambda and p-value determined for each tumour, with a median p-value<0.05 indicating significant phylogenetic signal for that gene. Genes with recurrent phylogenetic signal were defined as those with significant phylogenetic signal in at least 3 tumours. The STRINGdb R package (Szklarczyk et al., 2021) was used to determine pathway enrichment of these recurrent phylogenetic genes and “string-db.org” used for plotting PPAR signalling genes.

In the pathway phylogenetic signal analysis, pathway enrichment values were used as input for ‘phylosig’ for the 48 pathways. Significance was then determined as above. Recurrent phylogenetic pathways were defined as pathways with significant phylogenetic signal in at least 2 tumours and Fisher’s exact tests were used to determine enrichment/depletion in pathway groups and classes.

To determine the power for each tumour used in the phylogenetic signal analysis, gene expression was simulated and lambda p-values estimated. Gene expression was Poisson distributed across nodes and was increased by a factor of 5-100% across every clade of the tree. This was performed over the forest of 100 trees with differing branch lengths and this process was then repeated 1000 times. The power to detect significant phylogenetic signal for a particular expression % change at a particular clade was therefore inferred by the percentage of simulations which had a median (i.e. over the 100 branch length variant trees) p-value <0.05.

### Genetic determinants of gene expression heterogeneity

Tumours with at least two tumour samples were included in this analysis (153 tumour samples from 19 tumours, median 4 samples per tumour) and only loci mutated in at least two samples and connected to an expressed genes (gene Groups 1-3 from Figure 1) were analysed (22,961 mutated loci connected to 5,927 expressed genes – 29,949 unique gene-mutation combinations).

The following data was used as input for the linear model:

- Exp: A gene x sample matrix of the variance stabilised normalised gene expression of Group 1-3 genes converted into a z-score by minusing the mean expression of all samples and dividing by the standard deviation of all samples.
- CNA: A gene x sample matrix of the total copy number of the gene locus. If multiple copy number states were detected for the same gene, the segment which overlapped the most with the gene’s locus was selected.
- Mut: A binary mutation x sample matrix where mutations (SNVs and indels) were either within the enhancer region of the gene or a non-synonymous mutation within the coding region of the gene itself. Enhancer links to genes were defined using ‘double-elite’ annotations from GeneHancer tracks (Fishilevich et al., 2017). Some enhancer regions overlapped with the gene coding region and non-synonymous mutations in these regions were annotated as both enhancer and non-synonymous.
- Purity: The purity of each sample as determined from dWGS or sWGS.

In addition, 14 matched normal samples were added and these were assigned WT for all mutations, 2 for total copy number and 0 for purity. For each gene-mutation combination the following linear model was implemented:

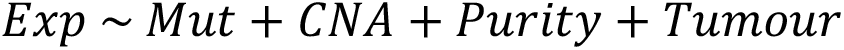

Where ‘Tumour’ indicated whether the sample was a normal or tumour sample.

A gene-mutation combination was said to be explained if the FDR adjusted p-value of the F-statistic for overall significance was less that 0.01. A gene-mutation combination was significantly affected by a variable (i.e. Mut/CNA/Purity/Tumour) if the FDR-adjusted p-value for the coefficient of that variable was <0.05.

For the analysis of clonality (Fig 5G), a gene was considered to be associated with an ‘eQTL’ if at least one mutation combination for that gene was significant for Mut while a gene was considered ‘subclonal’ if at least one mutation associated with that gene was not found in all matched DNA-RNA samples for at least one tumour. For combining eQTLs with the phylogenetic analysis and clonality (Fig 5H), a gene mutation combination was considered an ‘eQTL’ if it was significant for Mut, considered ‘subclonal’ if it was not found in all matched DNA-RNA samples for at least one tumour and considered ‘phylogenetic’ if the associated gene had significant phylogenetic signal in at least three tumours.

To look for recurrence of eQTL mutations in the Hartwig cohort, mutation loci were first converted to hg19 using liftOver from the rtracklayer R package (Lawrence et al., 2009) and the “hg38Tohg19.over.chain” from http://hgdownload.cse.ucsc.edu/goldenpath/hg38/liftOver. 2/22959 loci could not be converted and were therefore discarded for this analysis. The converted loci were searched for in the CRC Hartwig cohort using the “purple.somatic.vcf.gz” files. For the Hartwig gene expression, the ‘adjTPM’ values were used and converted into a z-score while tumour purity was extracted from the metadata. For each locus that had at least one mutated DNA-RNA Hartwig sample the linear models of Exp~Mut+Purity and Exp~Purity were compared via a likelihood ratio test. An eQTL was said to validate in Hartwig if the p-value of the test was <0.05 and the coefficient of the ‘Mut’ variable was the same sign as the coefficient in the original eQTL analysis (i.e. the mutation increased expression in EPICC and Hartwig or vice versa).

A *post hoc* power analysis was carried out using the pwr.t2n.test from the pwr R package. For each eQTL, the absolute mutation effect size was used as the input effect size with power set to 0.99 and n2 set to ‘the number of DNA-RNA Hartwig CRC samples (n=394) minus the number of Hartwig samples with the mutation’. The tool then returned the number of samples needed to see the effect and this number was multiplied by 1.15 given the non-parametric nature of the data. Note that if the absolute input effect size was greater than 3.04, this was set to 3.04 since higher values returned an NA result.

### Combining with ATAC-seq

Enhancer eQTLs that were subclonal in at least one tumour according to matched DNA-RNA data were investigated with matched ATAC-seq data (processing of ATAC-seq data detailed in methods of epigenome paper). For each mutation in each tumour, samples with the mutation were plotted as a separate track to wild-type samples for the enhancer region and mutation locus indicated. A t-test was performed and log-fold change calculated based on the mean normalised read counts for mutant and wild-type samples. An eQTL was said to be associated with a change in ATAC if the p-value for the test was less than 0.05.

## Acknowledgments

This study was principally supported by funding from the Wellcome Trust (202778/Z/16/Z to T.A.G. and 202778/B/16/Z to A.S.) and the Medical Research Council (MR/P000789/1 to A.S.). A.S. and T.A.G. were also supported by Cancer Research UK (A22909 and A19771) and the National Institute of Health (NCI U54 CA217376 to D.S., T.A.G. and A.S. This work was also supported by a Wellcome Trust award to the Centre for Evolution and Cancer at the ICR (105104/Z/14/Z). D.R. was partially supported by a Bicocca 2020 Starting Grant and by a Premio Giovani Talenti dell’Università degli Studi di Milano-Bicocca. L.M. is supported by Cancer Research UK (A23110).

## Data availability

Analysed data are available on Mendeley: https://data.mendeley.com/datasets/dvv6kf856g/2. Sequence data have been deposited at the European Genome-phenome Archive (EGA), which is hosted by the EBI and the CRG, under accession number EGAS00001005230. Further information about EGA can be found on https://ega-archive.org.

## Code availability

Complete scripts to replicate all bioinformatic analysis and perform simulations and inference are available at: https://github.com/sottorivalab/EPICC2021_data_analysis.

## Author contributions

J.H. analysed and interpreted the data, with focus on RNA data. T.H. analysed and interpreted the data, with focus on ATAC and WGS data. GDC performed copy number analysis. C.L. performed ATACseq analysis. I.S. devise multi-omics protocol, collected the samples and generated the data. C.K. collected the samples and contributed to data generation. C.L. contributed to ATAC data analysis. M.M., J.F.M., A.M.B., contributed to data generation. B.J. analysed methylation array data. L.Z.O. contributed to dN/dS data analysis. C.J., E.L., G.Car. contributed to data analysis. D.N. and K.C. contributed to signature analysis. A.B. generated methylation array data. I.B. contributed to analysis of CTCF binding site mutations. H.C., M.M., and M.J. contributed to tissue collection. D.S. contributed to experimental design and data interpretation. J.B. contributed to sample collection coordination. M.R.J. supervised sample collection. L.M. contributed to result interpretation. T.A.G. and A.S. conceived and supervised the study and wrote the manuscript.

