## Supplementary Figures for "Phenotypic plasticity and genetic control in colorectal cancer evolution"

### Supplementary Figure 1

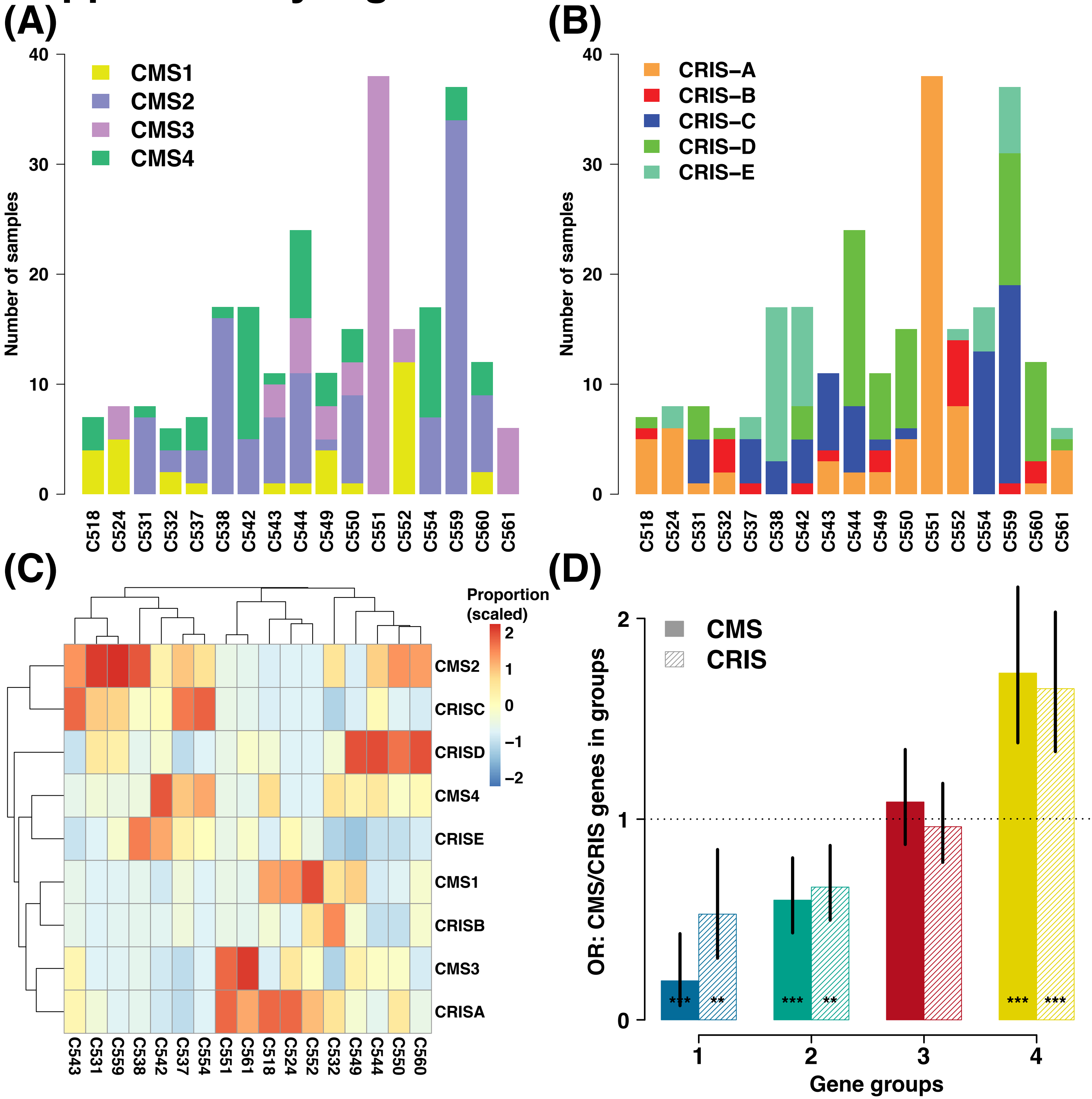


**Figure S1.** CMS and CRIS classification heterogeneity. **(A)** Classification of CMS in multi-region samples. **(B)** Classification of CRIS in mult-region samples. **(C)** Proportional heatmap based on data in (A) & (B) showing associations between particular CMS and CRIS classes. **(D)** Results of Fisher's exact tests showing that both CMS and CRIS template genes are depleted in gene Groups 1&2 but are enriched in Group 4 genes.

### Supplementary Figure 2 A-D

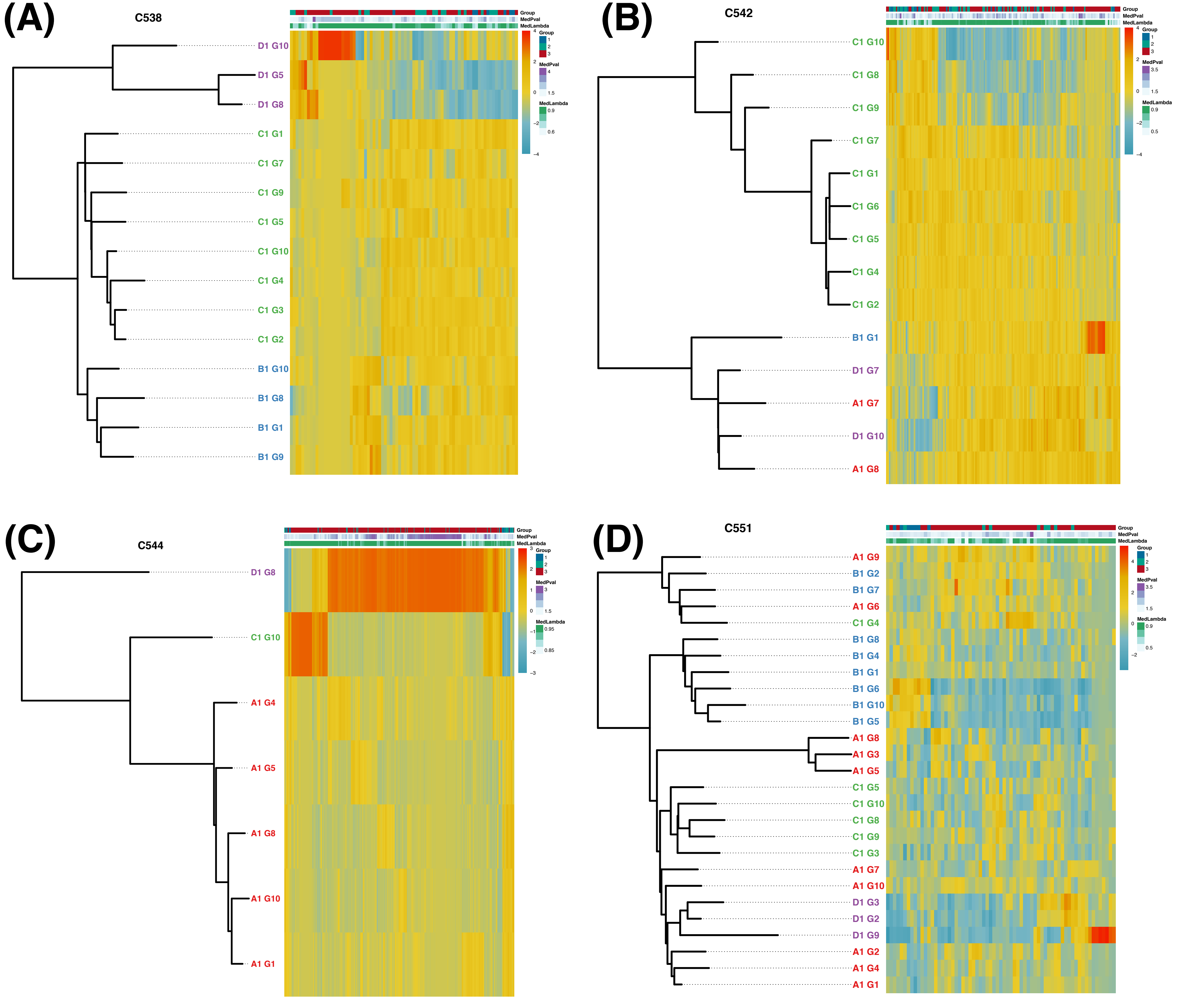

### Supplementary Figure 2 E-H

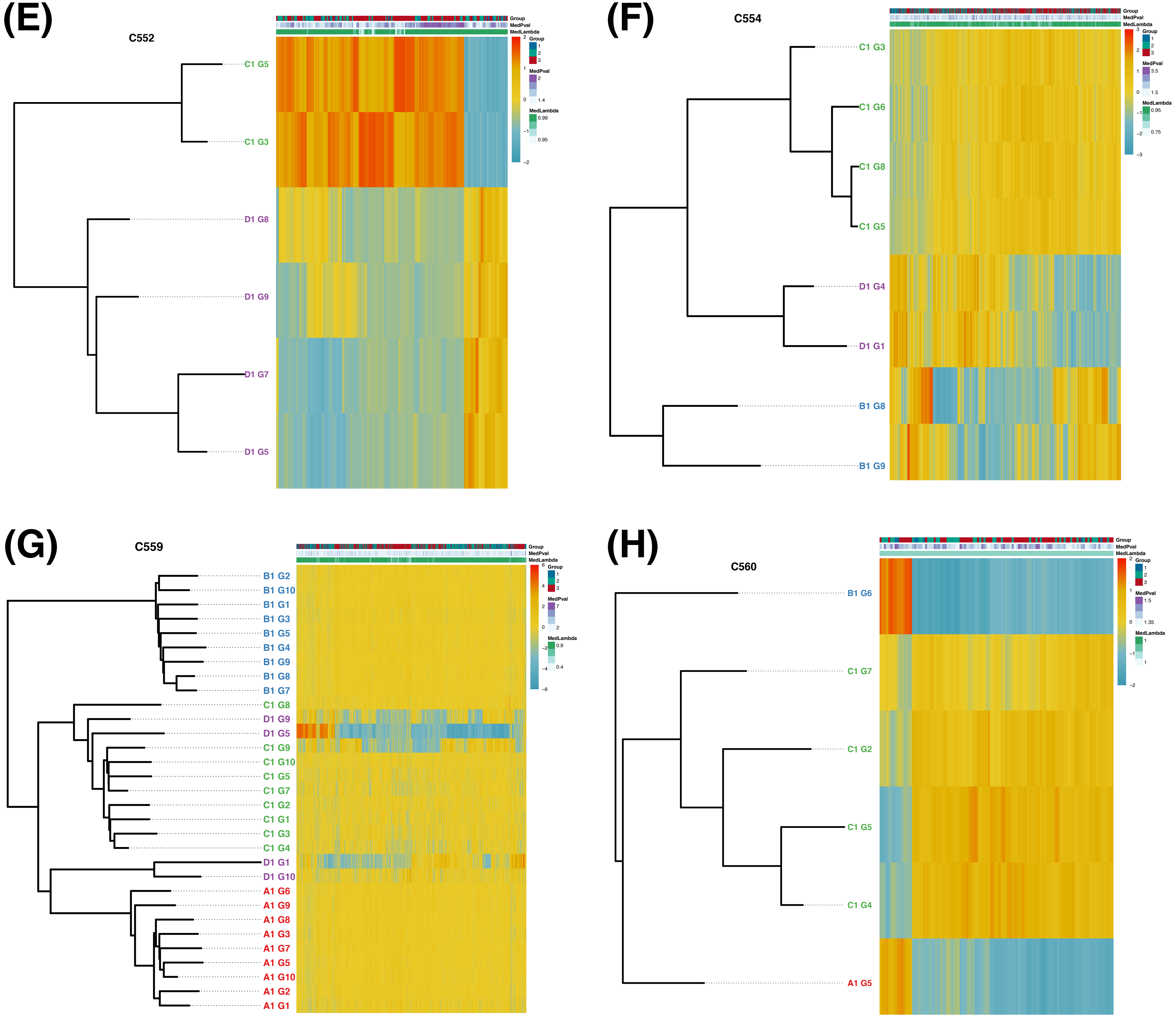


**Figure S2.** Phylogenetic trees annotated with the expression of genes which had significant phylogenetic signal.

### Supplementary Figure 3

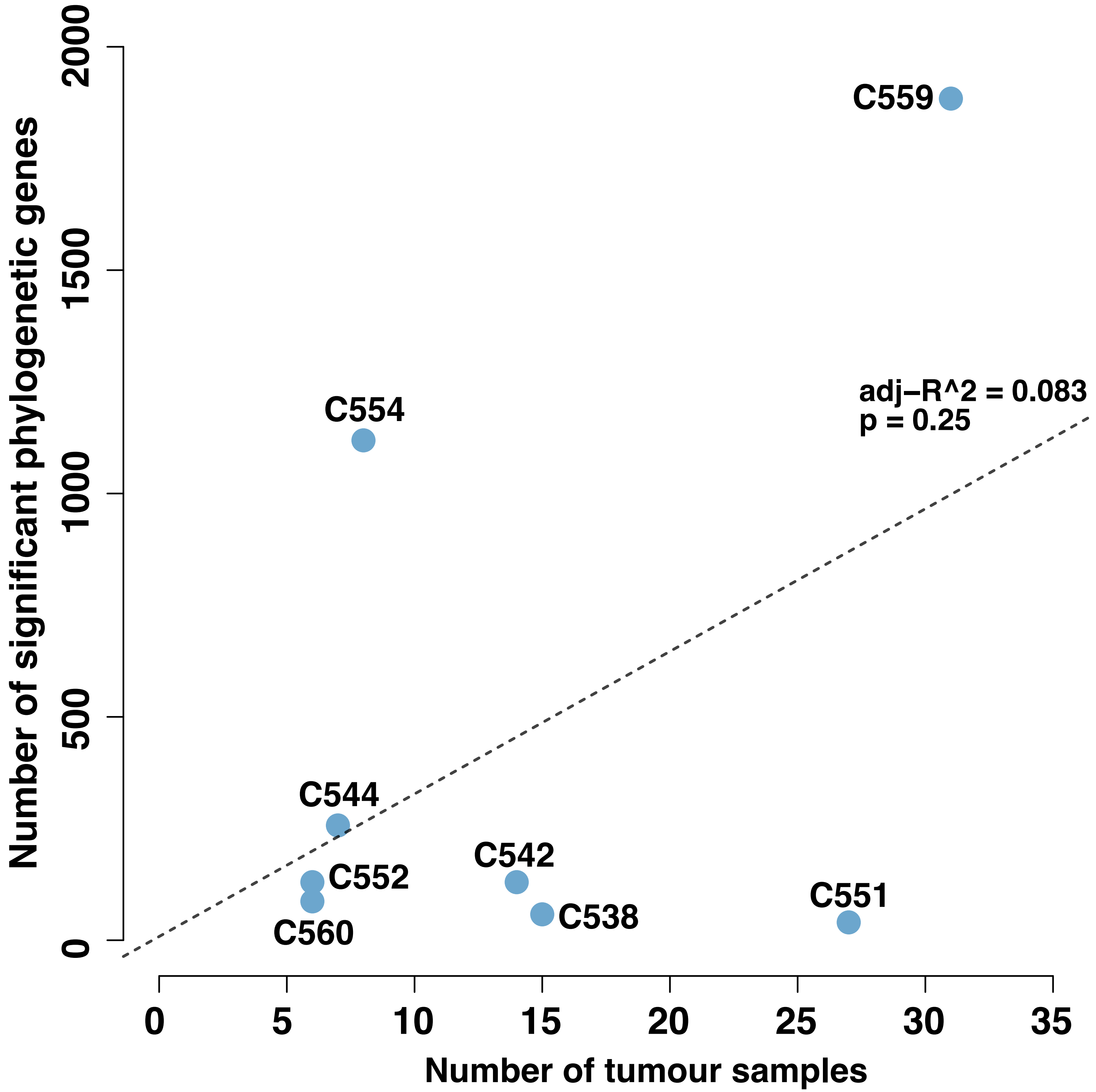


**Figure S3.** Correlation (or lack thereof) of the number of significant of phylogenetic genes per tumour against the number of tumour samples analysed.

### Supplementary Figure 4 A&B

(A)

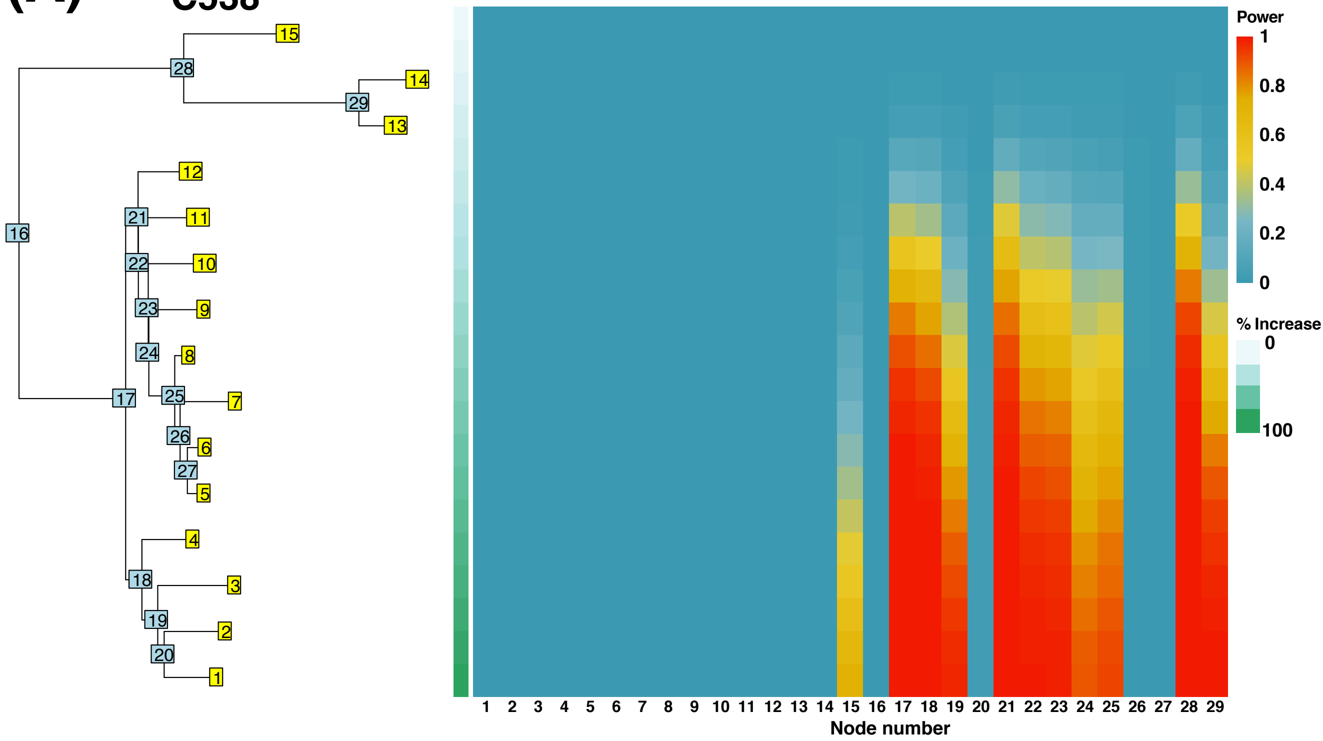


(B)

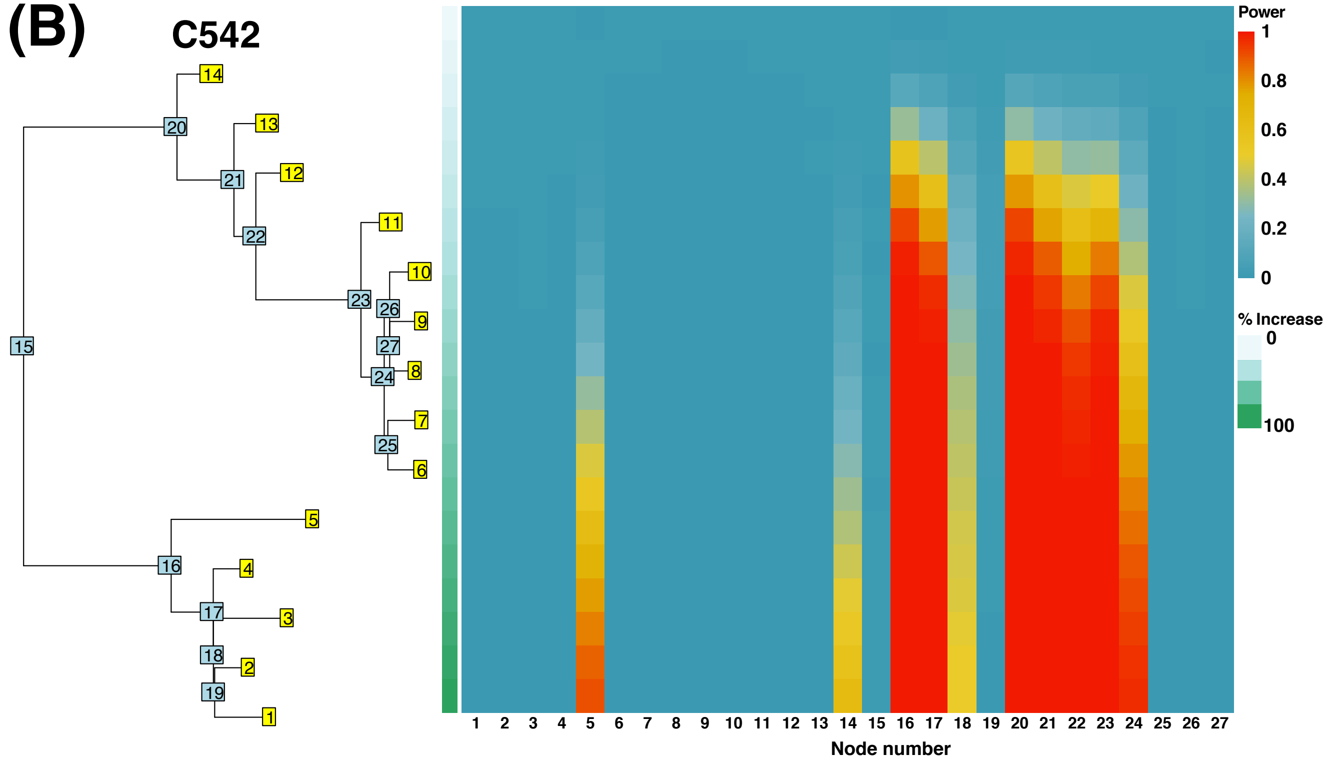

### Supplementary Figure 4 C&D

(C)

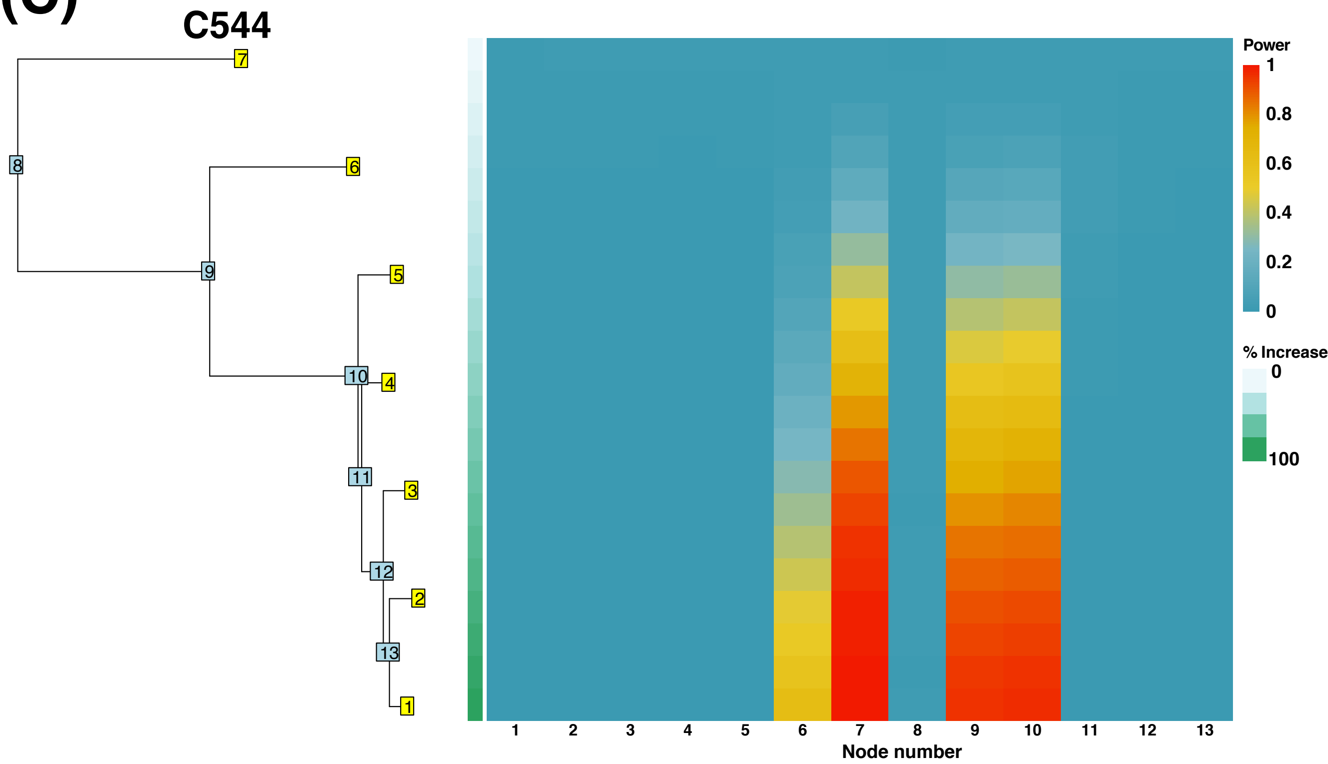


(D)

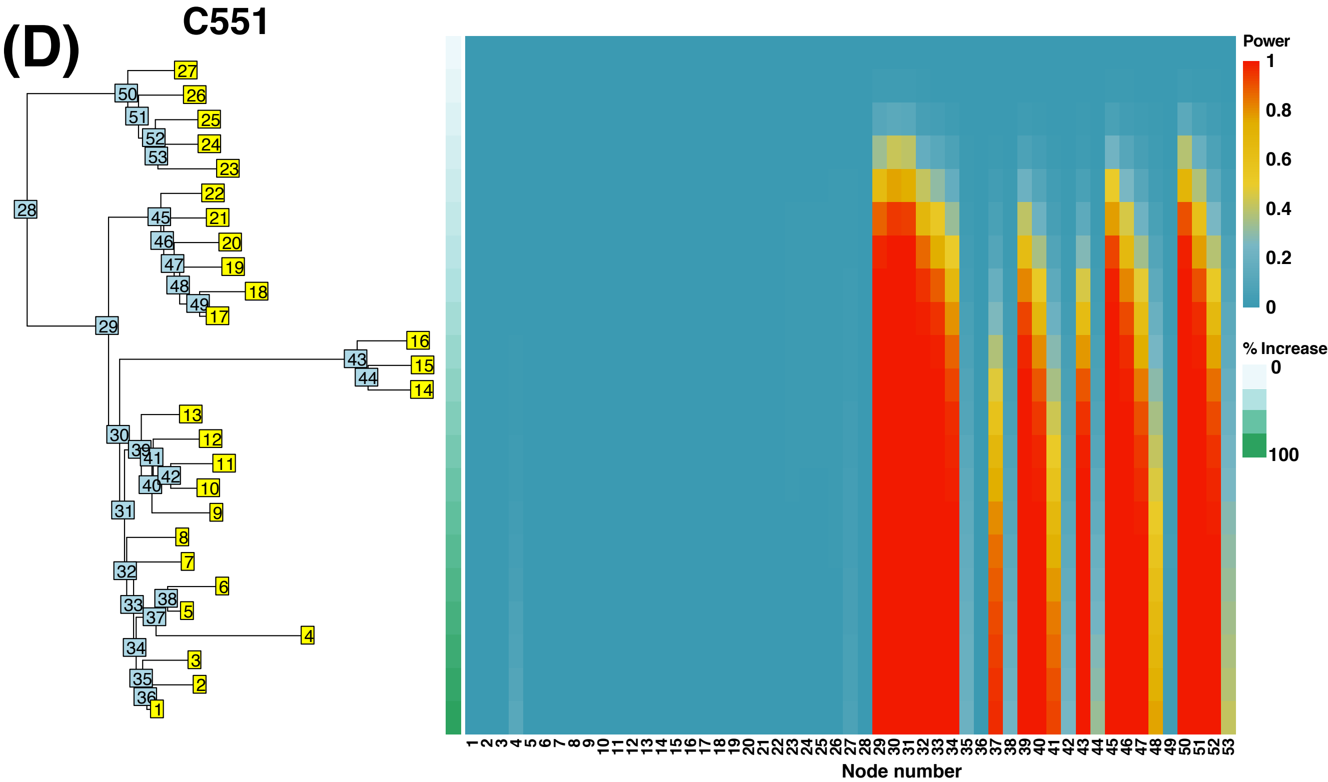

### Supplementary Figure 4 E&F

(E)

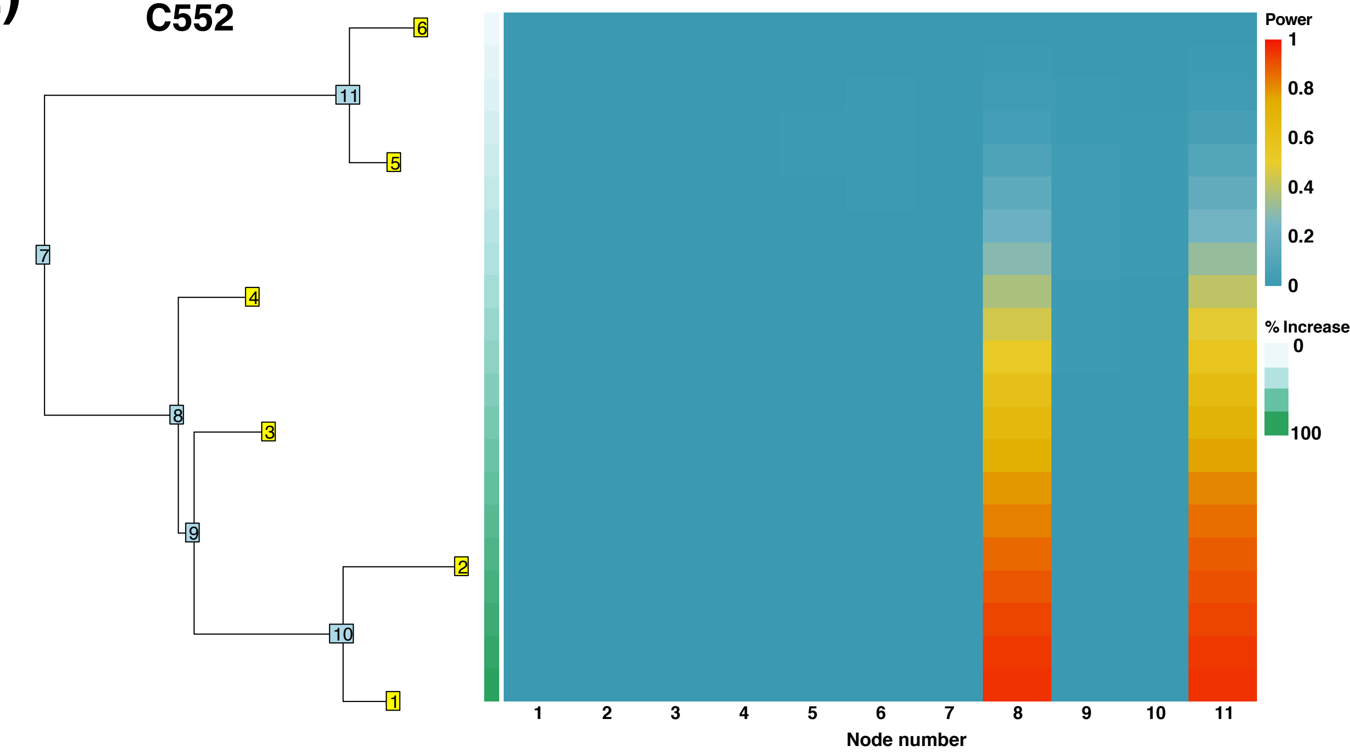


(F)

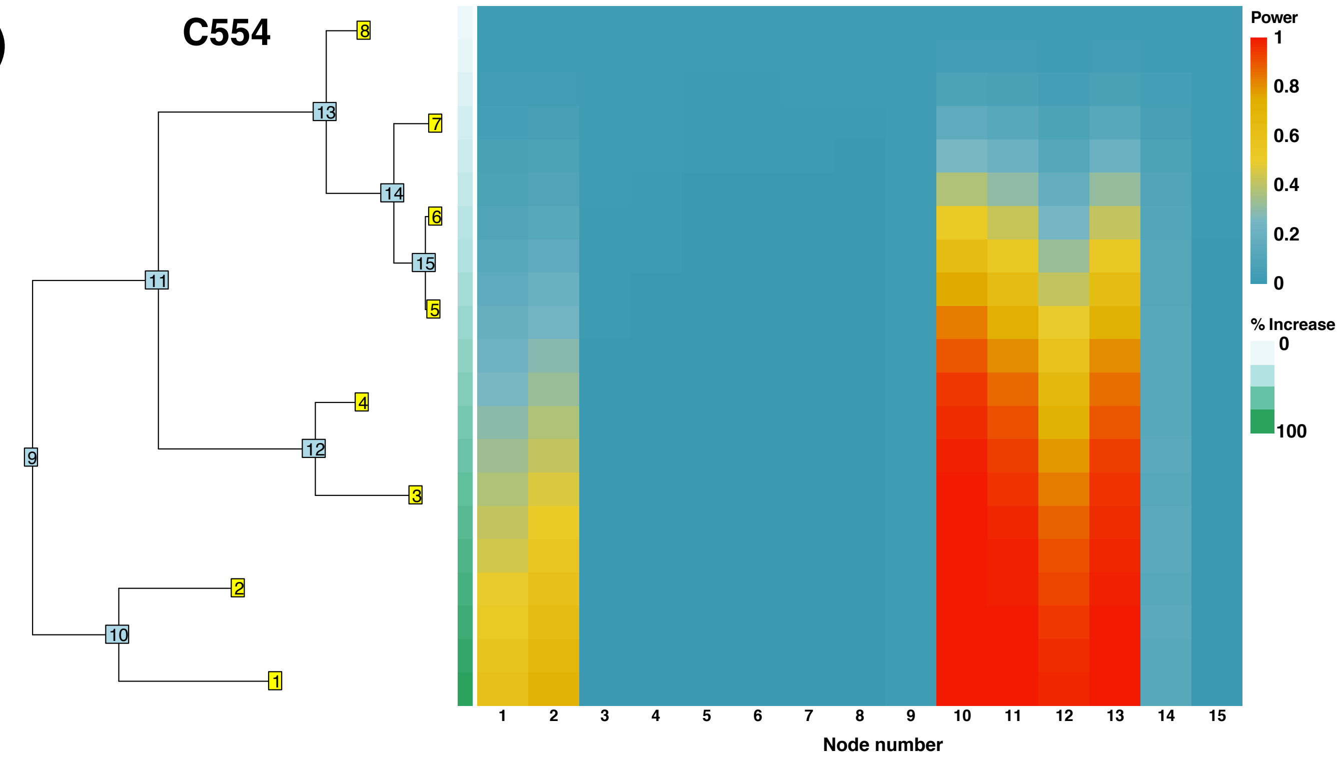

### Supplementary Figure 4 G&H

(G) C559

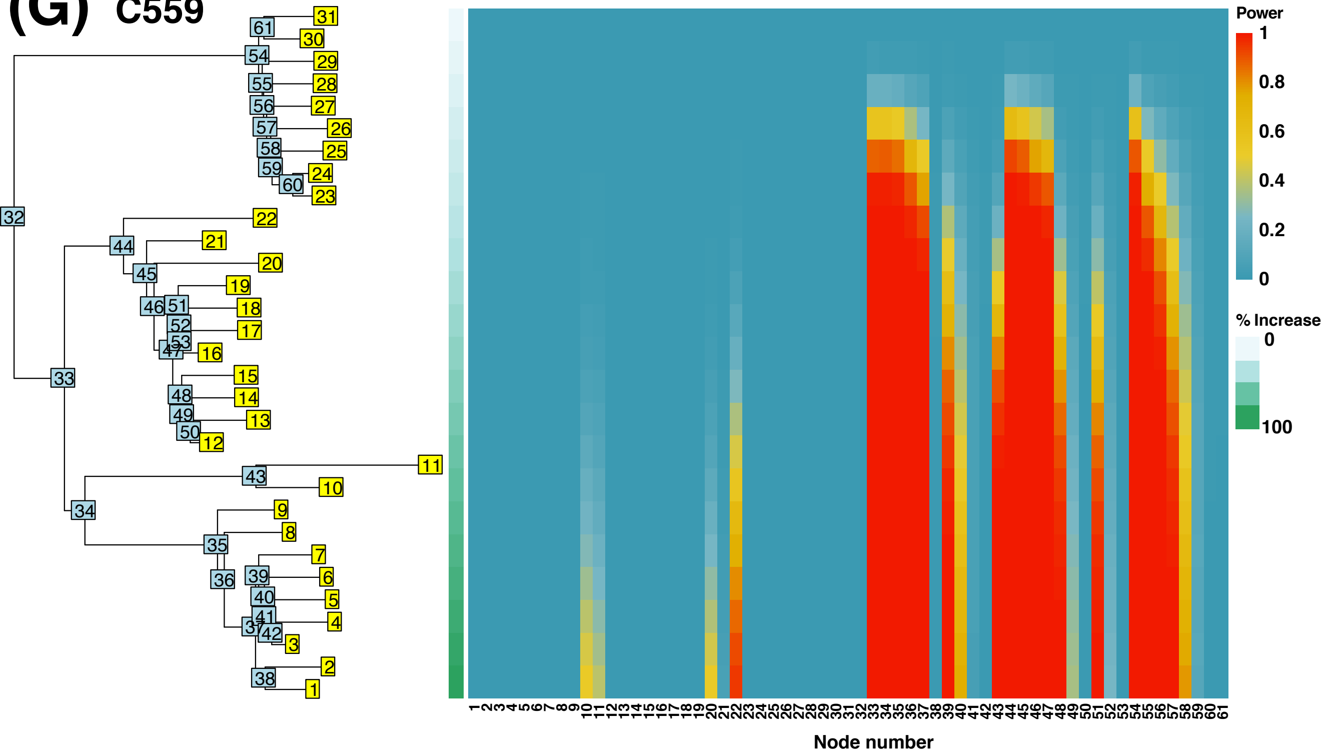


(H) C560

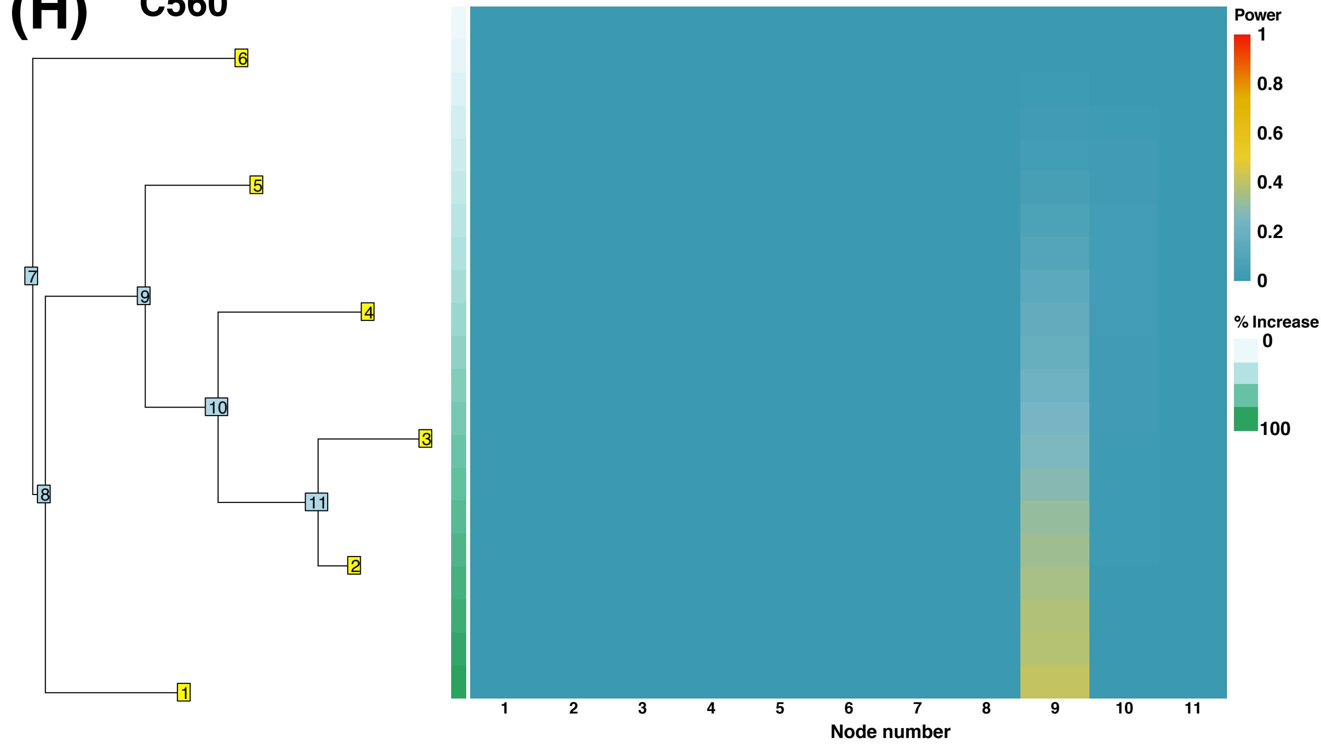


**Figure S4.** Analyses to detecdetermine the power to detect phylogenetic signal for particular nodes and certain percentage expression changes for multi-region tumours.

### Supplementary Figure 5 A-D

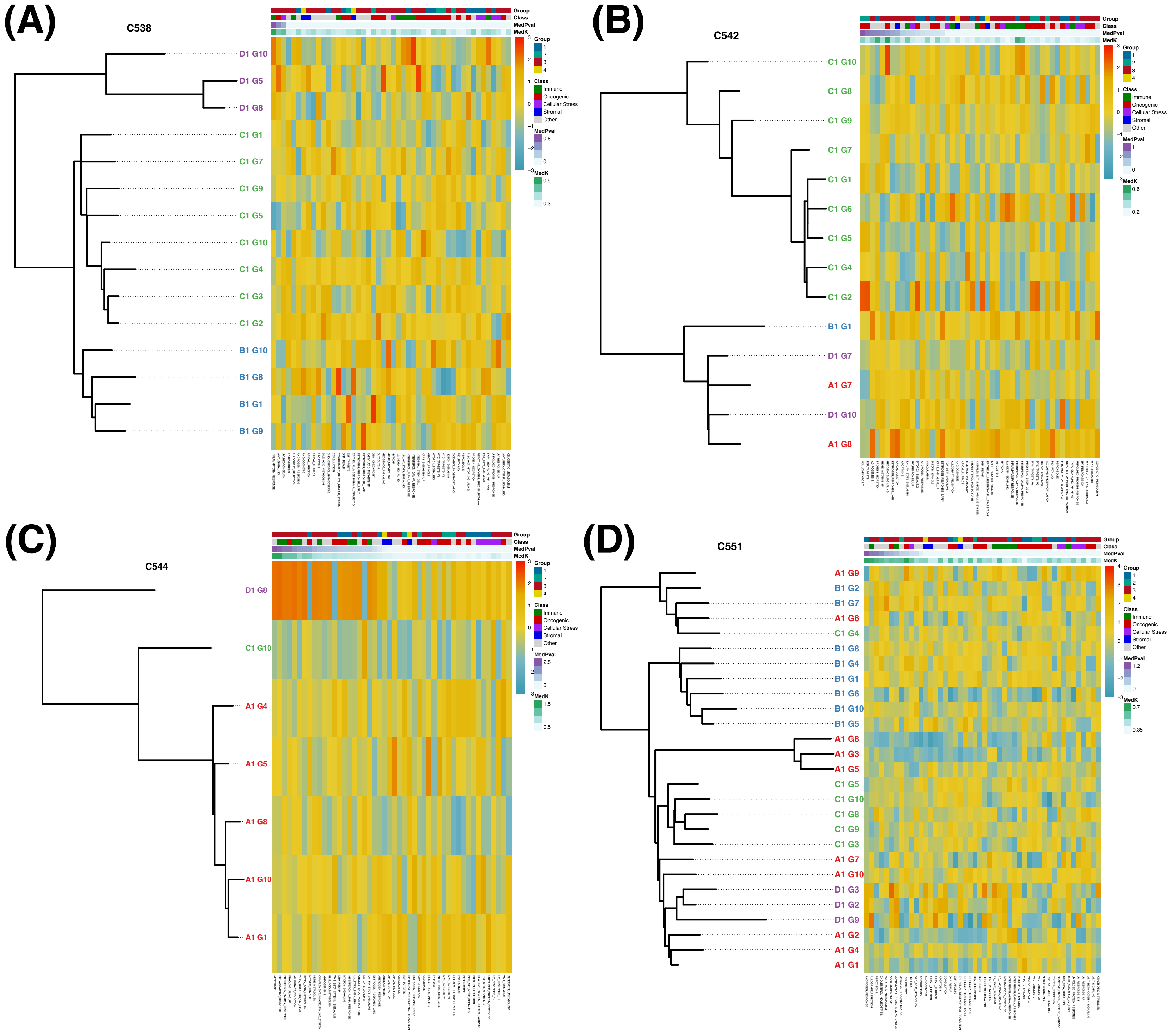

### Supplementary Figure 5 E-H

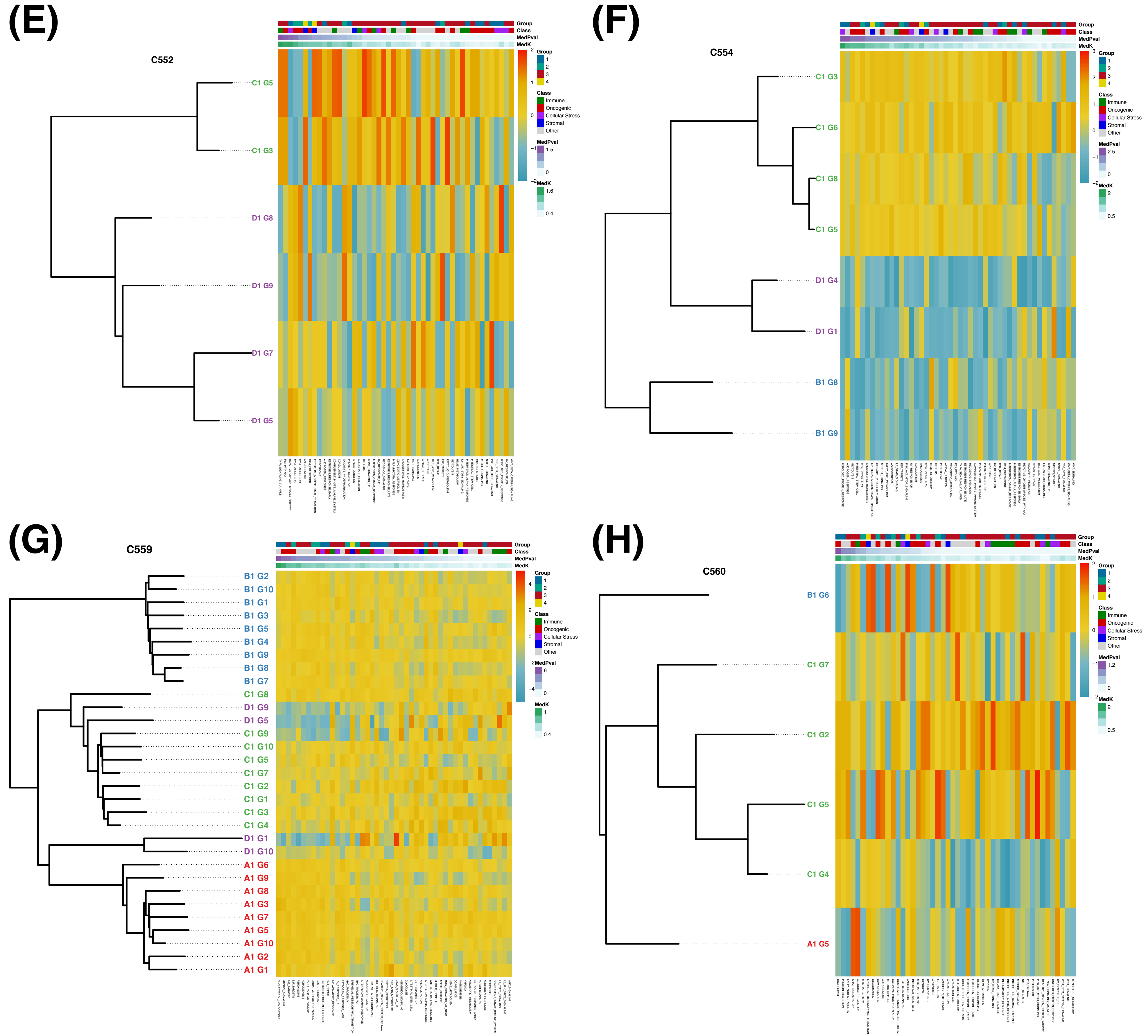


**Figure S5.** Phylogenetic trees annotated with pathway enrichment scores.

### Supplementary Figure 6

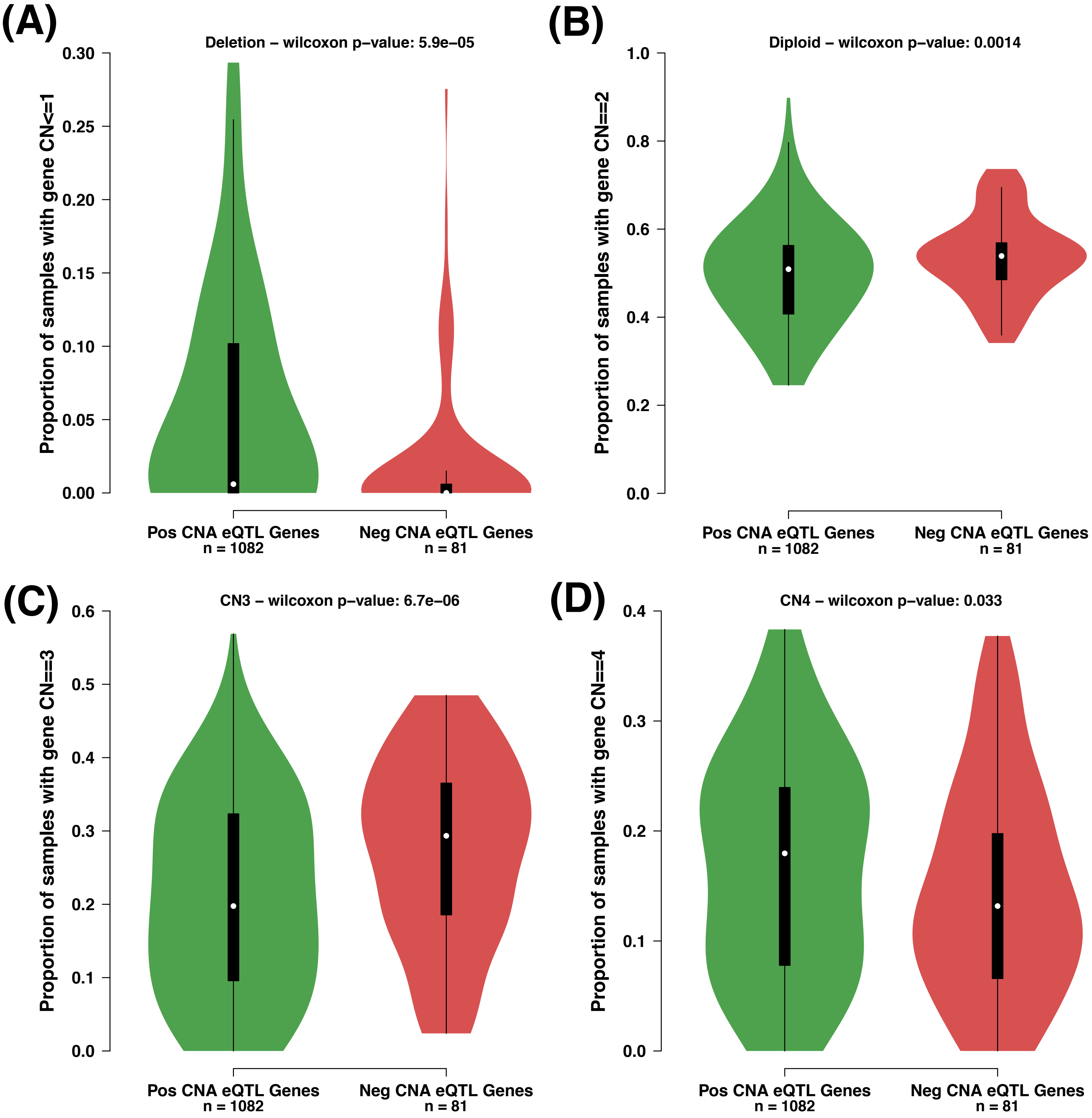


**Figure S6.** CNA positive vs negative copy number associations

### Supplementary Figure 7 A-I

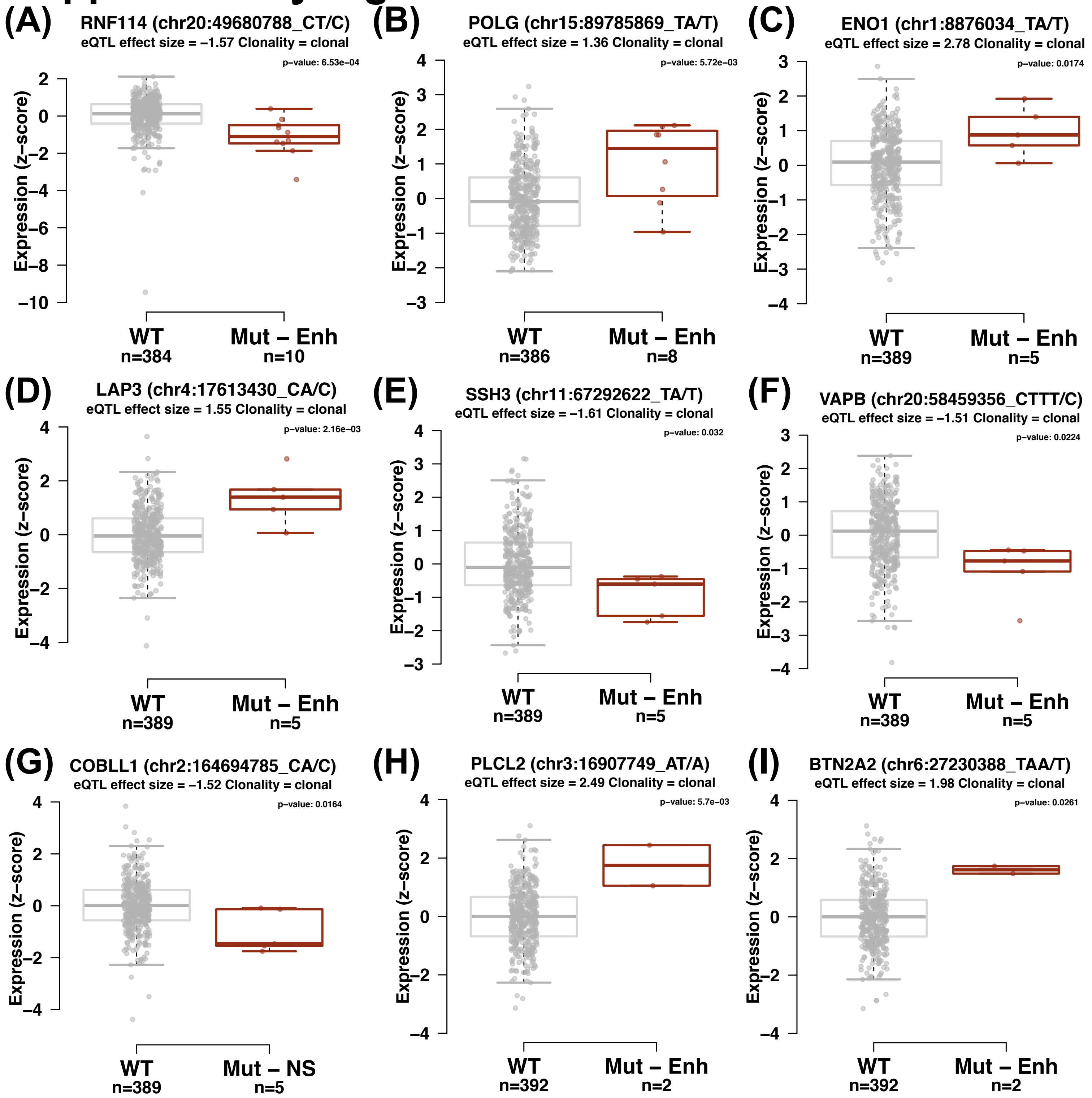

### Supplementary Figure 7 J

(J)

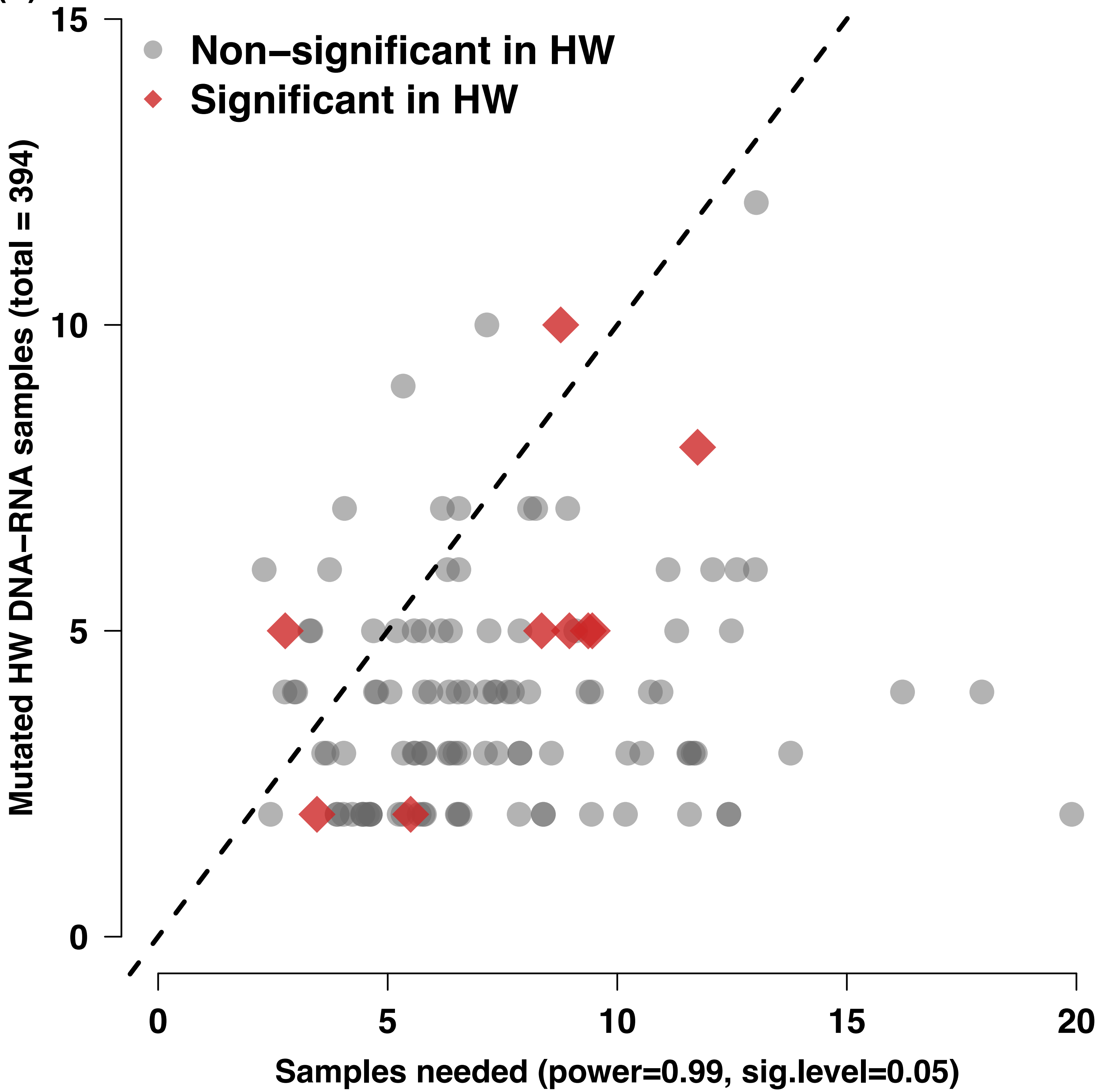


**Figure S7.** Investigating eQTLs in the Hartwig mCRC cohort. **(A)-(I)** Recurrent eQTLs in the Hartwig mCRC which also significantly correlate with expression changes in mutated samples. **(J)** Power analysis demonstrating the lack of power to detect most eQTLs in the Hartwig cohort.

### Supplementary Figure 8

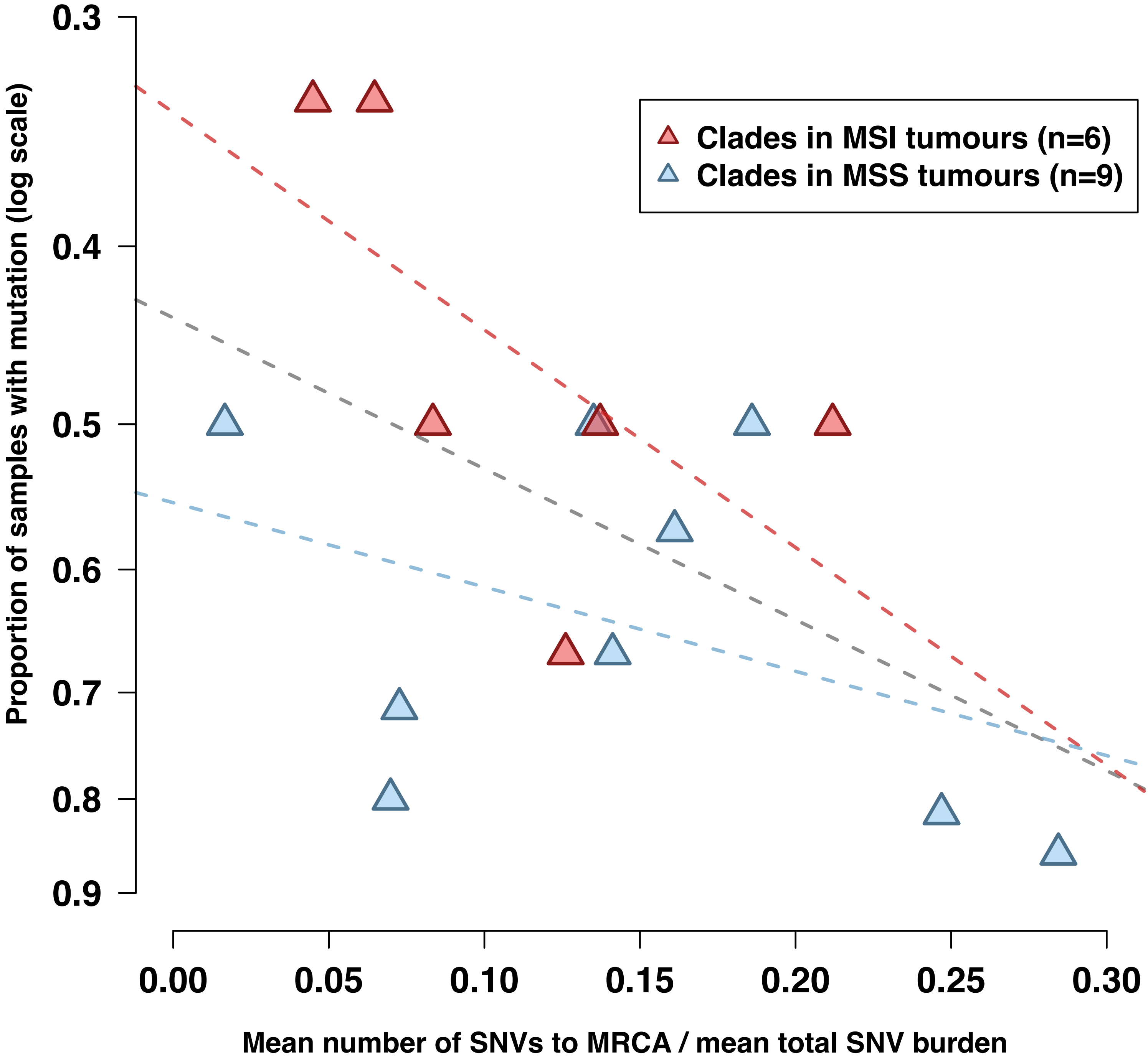


**Figure S8.** The time to most recent common ancestor (MRCA; measured in proxy by mean number of SNVs to MRCA divided by mean total SNV burden) correlates with clone size (-ln proportion of samples with mutation) for clades mutated with eQTLs.

### Supplementary Figure 9 A-D

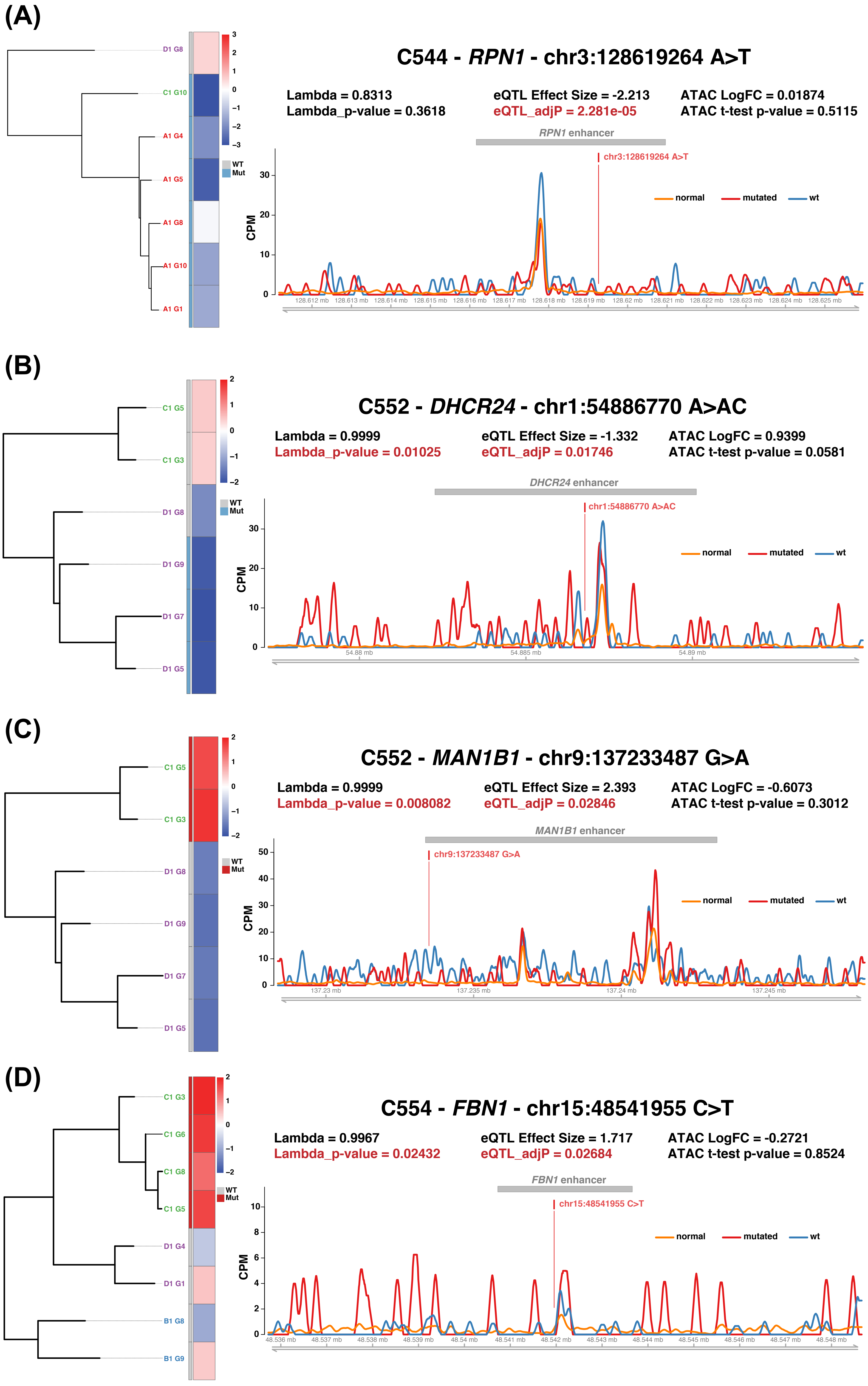

### Supplementary Figure 9 E-G

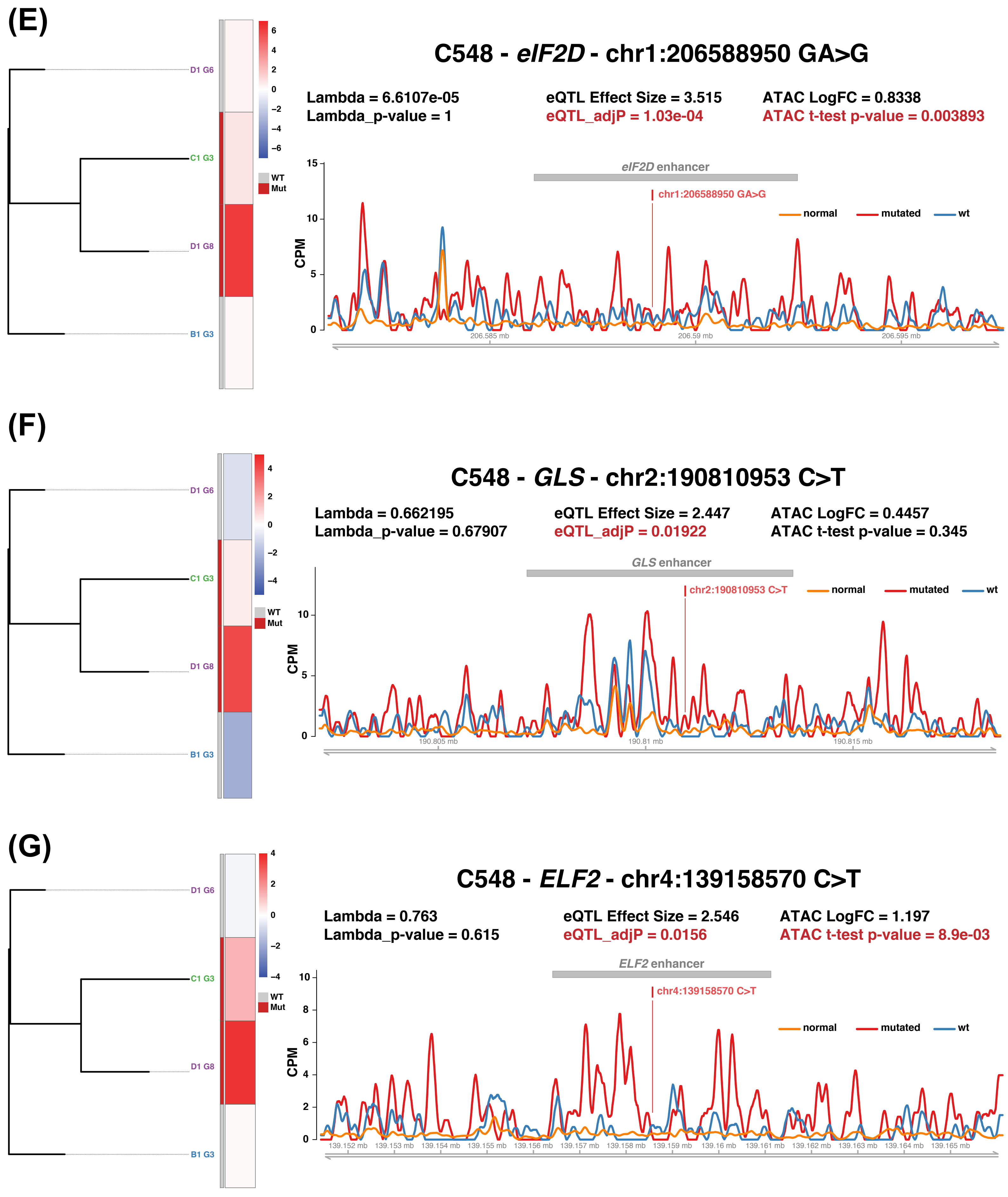

### Supplementary Figure 9 H-J

(H)

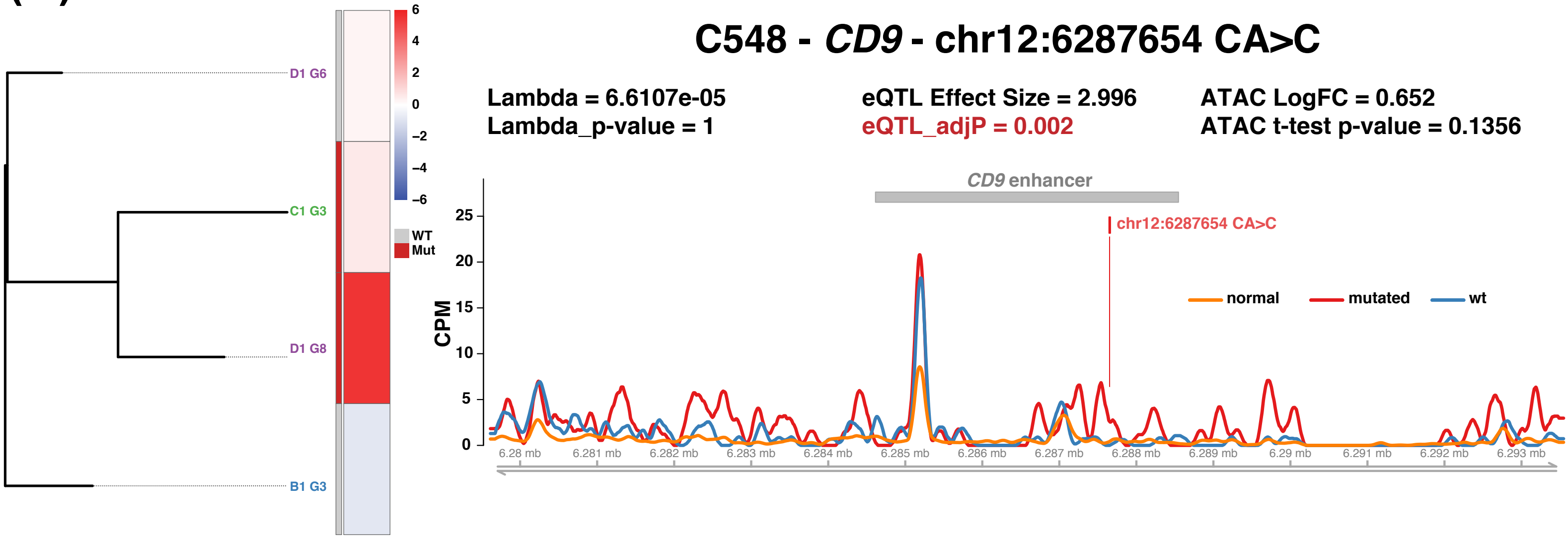


(I)

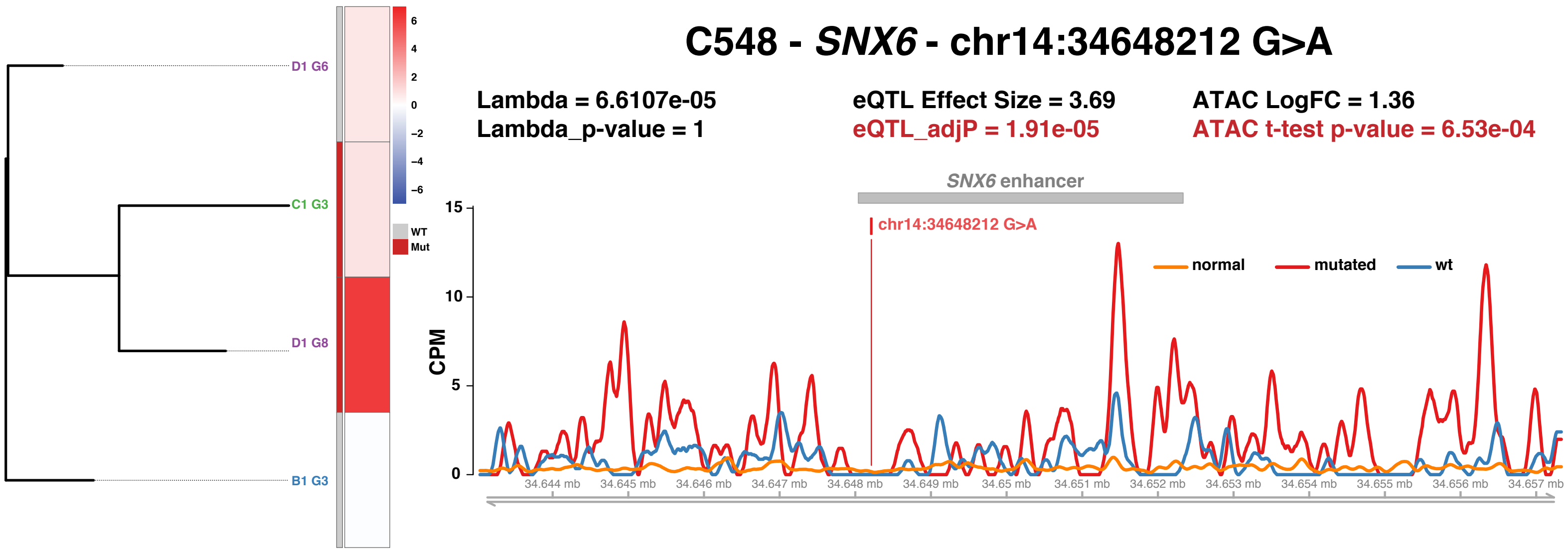


(J)

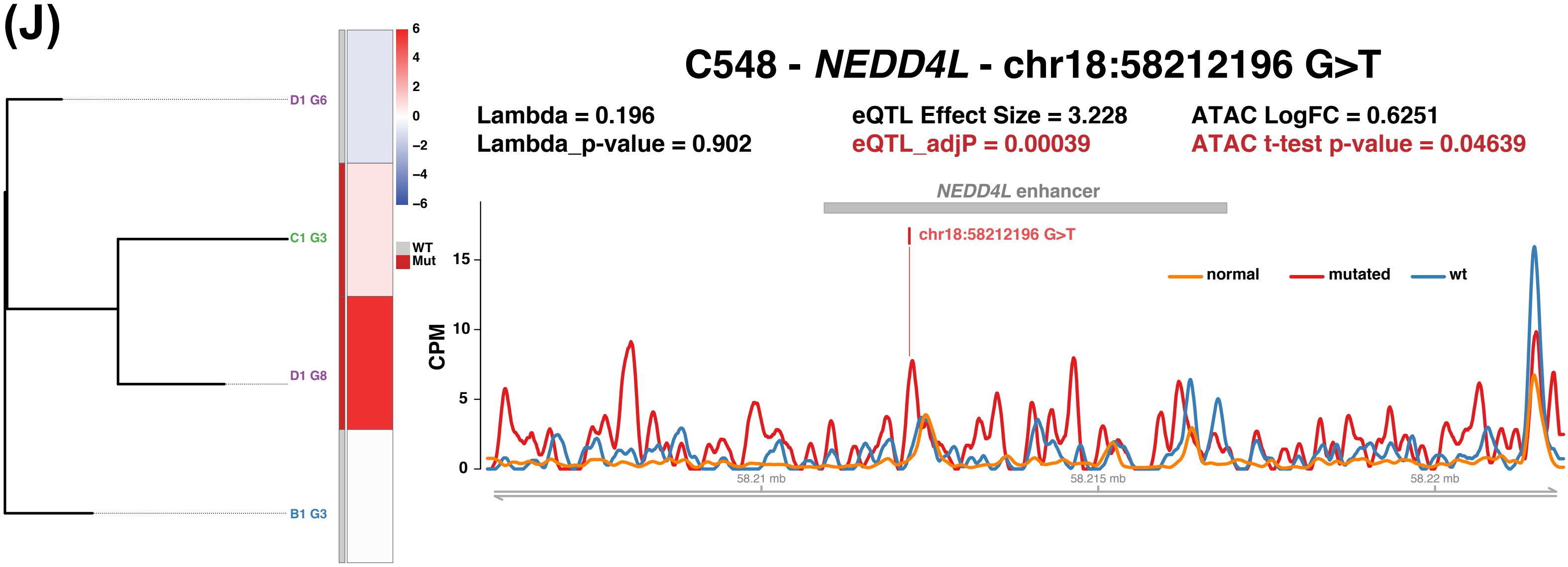


**Figure S9.** Combining eQTLs ATAC-seq and phylogenetic data. Example sub-clonal eQTLs with supporting ATAC-seq data from tumours (A) C544, (B&C) C552, (D) C554 and (E-J) C548.
